# Emergence of new enhancers at late DNA replicating regions

**DOI:** 10.1101/2022.12.22.521323

**Authors:** Paola Cornejo-Páramo, Veronika Petrova, Xuan Zhang, Robert S. Young, Emily S. Wong

## Abstract

Enhancers are fast-evolving genomic sequences that control spatiotemporal gene expression patterns. By examining enhancer turnover across mammalian species and in multiple tissue types, we uncovered a relationship between the emergence of novel enhancers and genome organization as a function of germline DNA replication time. While enhancers are most abundant in euchromatic regions, new enhancers emerged almost twice as often in late compared to early germline replicating regions, independent of transposable elements. Using a sequence model, we demonstrate that new enhancers are enriched for mutations that alter transcription factor (TF) binding. Recently evolved enhancers appeared to be mostly neutrally evolving and enriched in eQTLs. They also show more tissue specificity than conserved enhancers, and the TFs that bind to these elements, as inferred by binding sequences, also show increased tissue-specific gene expression. We find a similar relationship with DNA replication time in cancer, suggesting that these observations may be time-invariant principles of genome evolution. Our work underscores that genome organization has a profound impact in shaping mammalian gene regulation.

## Introduction

Enhancers are *cis*-regulatory elements essential as modulators of spatiotemporal gene expression by acting as integrators of trans-acting signals by recruiting transcription factors (TFs) and other effector molecules. Enhancers are typically rapidly evolving and are frequently species-specific ^1–4^. For example, most human enhancers are not found in the mouse ^3^.

The factors responsible for enhancer turnover are not well understood. The prevailing model of enhancer evolution is the mobilization of transposable elements (TE) and their insertions to new genomic locations ^5^. TEs often overlap enhancer elements, and thus, they have been hypothesized to play a major role in the dynamic landscape of enhancer turnover in mammals by distributing cis-regulatory elements across the genome. However, they do not account for most recently evolved mammalian enhancers, many of which appear to have originated from ancestral sequences without prior biochemical activity in the same tissue ^6–10^.

Notably, local point mutations in non-regulatory sequences can give rise to novel enhancer activity, suggesting a possible mechanism for generating tissue-specific enhancers ^11–16^. DNA replication time is one of the most significant predictors of local mutational density ^17–19^. *De novo* mutations are elevated towards the latter stages of the S-phase. This phenomenon is likely due to errors made in DNA replication and a reduced ability of DNA repair mechanisms to function effectively during late replication time ^20^. In hominids and rodents, mutation rates are 20-30% higher at late compared to early replication domains ^21^. The trend of increased mutational burden during DNA replication extends throughout eukaryotic evolution ^22^. The DNA replication timing program, defined by the temporal order of DNA replication during the S-phase, is also closely linked to the spatial organization of chromatin in the nucleus and transcriptional activity ^23^. Late replicating domains are linked to facultative heterochromatin and tissue-specific gene expression ^24,25^.

Thus, we hypothesized that DNA replication timing plays a role in the emergence and diversification of enhancers through de novo mutations ^2,26,27^. In the context of enhancer turnover in mammals, we investigate the role of nucleotide substitutions linked to DNA replication timing. Using detailed maps of candidate cis-regulatory elements across species, we take a multi-scale approach to explore the relations between enhancer turnover and the genome. We examine the contribution of TEs and the *de novo* creation of TF binding sites to enhancer turnover. By comparing enhancers across DNA replication domains and their tissue-specific activity across vastly different timescales, we aim to illuminate the evolutionary trajectories of enhancers and their implications for gene regulation.

## Results

### Germline replication time is associated with the rate of enhancer turnover across the genome

Genetic changes occurring in the germline provide novel genetic variation that is the substrate for species evolution. We examined multi-tissue enhancer turnover in mice comparing across germline DNA replication time. Evolutionarily conserved and recently evolved, i.e., lineage-specific mouse enhancers were annotated using histone mark ChIP-seq data based on multi-species comparisons (cat, dog, horse, macaque, marmoset, opossum, pig, rabbit, rat) ^9^. Following convention, candidate enhancers are defined as sequences enriched for H3K27ac but absent in H3K4me3 (termed “active”) or enriched for H3K4me1 (termed “poised”) ^9,28^ (**Fig. 1A-B**). To ensure robustness, all enhancers were identified using consensus regions defined by overlapping multiple biological replicates by a minimum of 50% of their length ^9^.

**Figure 1.**
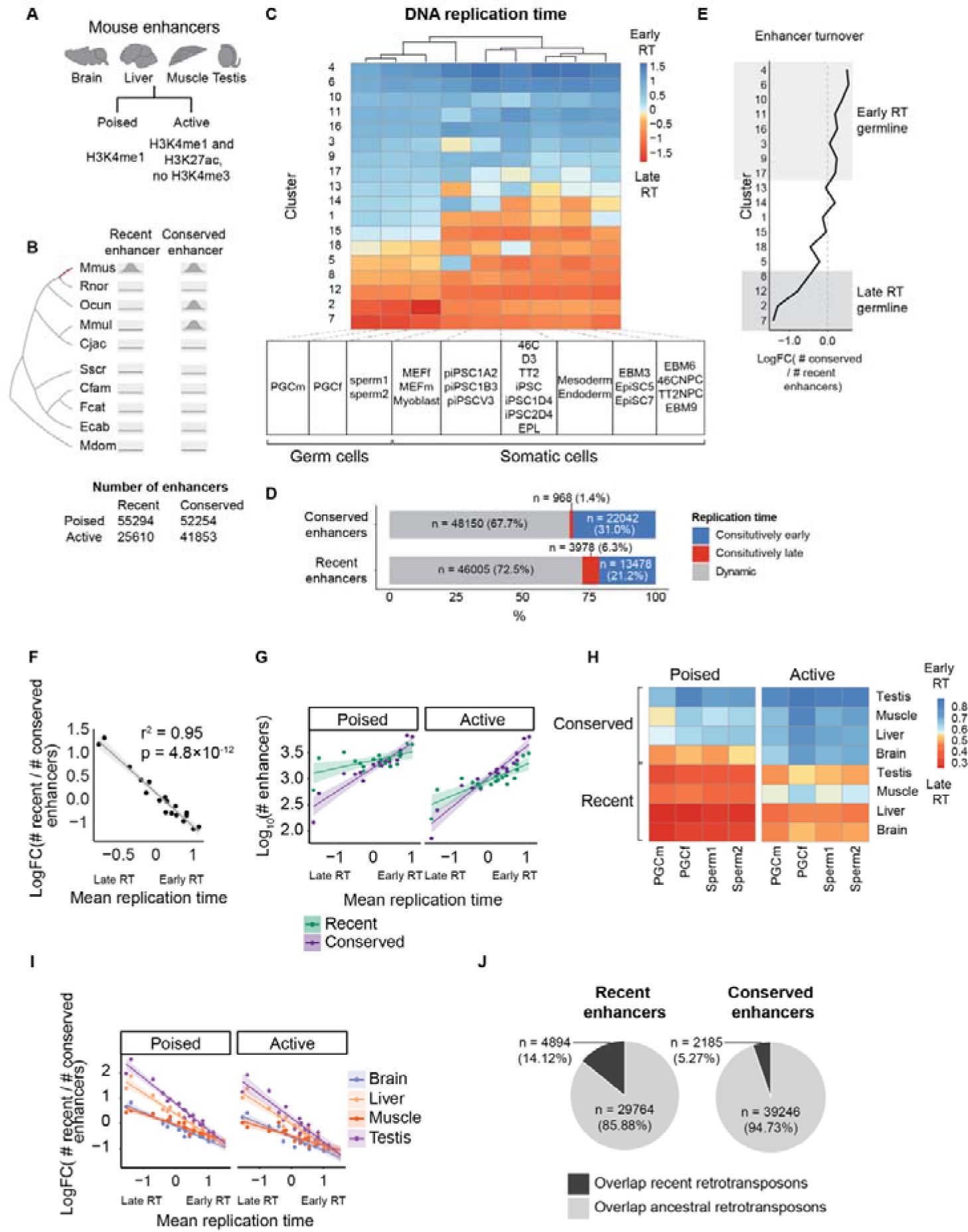
Enhancer turnover is coupled to germline replication timing. **(A)**. Mouse enhancers are defined based on combinations of histone marks. **(B)** Definition of mouse recent and conserved enhancers ^9^. Recent enhancers were defined as regions not aligning with nine other mammalian genomes in the same tissue ^9^. Conserved enhancers are aligned to regions with regulatory activity in at least two other species. Mmus = *Mus musculus*, Rnor = *Rattus norvegicus*, Ocun = *Oryctolagus cuniculus*, Mmul = *Macaca mulatta*, Cjac = *Callithrix jacchus*, Sscr = *Sus scrofa*, Cfam = *Canis familiaris*, Fcat = *Felis catus*, Ecab = *Equus caballus* and Mdom = *Monodelphis domestica*. **(C)** Replication time across 200 kb blocks of the mouse genome (n = 8966 blocks) in PGC (n = 2 cell lines), SSC cells (n = 2 cell lines), and early somatic cell types (n = 22 cell lines). Row clustering (blocks) was carried out with *k*-means clustering; columns are cell-type clusters generated with hierarchical clustering. Row clusters were ordered by mean DNA replication time, from early (top) to late (bottom), across columns (cell type clusters). **(D)** Number and percentage of recent and conserved enhancers in regions of **(C)** with constitutively early (blue), constitutively late (red) and dynamic (gray) replication time. Enhancers overlapping different replication time were not considered. **(E)** Enhancer turnover is the log fold change of conserved enhancers vs. recent enhancers for the 200 kb clusters across mean germline replication time calculated across PGC (n = 2) and SSP cell lines (n = 2). **(F)** Scatterplot of mean germline replication time (PGC + SSP) across the 18 clusters shown in **(C)**. R^2^ and p-value are indicated. **(G)** Scatterplot of germline mean DNA replication time (PGC + SSP) and log_10_-transformed numbers of recent and conserved enhancers. Each data point represents a cluster as defined in **(C)**. The shaded region represents the 95% confidence interval of the line of best fit. **(H)** Heatmaps of mean PGC and SSC DNA replication time of poised and active mouse enhancers separated by tissue and type. **(I)** Scatterplots of germline mean DNA replication time (PGC + SSP) and enhancer turnover by tissue and enhancer type. Each data instance corresponds to a cluster in **(C)**. The shaded region represents the 95% confidence interval of the line of best fit. **(J)** Pie charts of the number of recent/conserved enhancers overlapping recent/ancestral retrotransposons (Fisher’s exact test, p < 2.2 × 10^−16^, odds ratio = 2.99).

Enhancers were annotated as evolutionarily conserved if they possess enhancer-associated histone marks in at least two other species (n = 94107). Recently evolved enhancers were defined as cis-regulatory elements identified only in mice (n = 80904), where most of these regions aligned to non-regulatory regions in the genome of other species used in our comparisons (∼89%; liftOver -minMatch = 0.6). This supports similar findings in human enhancers ^3^. Both conserved, and species-specific enhancers showed a similar propensity to overlap ATAC-seq peaks, indicating comparable levels of chromatin accessibility (**Methods**, **Supplemental Table S1**).

To compare DNA replication timing, we obtained Repli-Seq data across the mouse genome from two germline stages: primordial germ cells (PGC) (n = 2, male and female) and spermatogonia stem cells (SSC) (n = 2), in addition to 22 other independent mouse cell lines across ten early stages of embryogenesis ^29,30^. Repli-seq resolves early and late replicating DNA by labeling it with nucleotide analogs ^31^. We assessed DNA replication time dynamics across cell types by partitioning the mouse genome into 200 kb regions and performing *k*-means clustering of cell-type specific DNA replication timing data across cell types (n = 8966 blocks, **Methods)**. This revealed approximately a third of the mouse genome to be consistently early or late replicating across both germline and developmental cell types, where 14% of the genome replicated early and 19% are late replicating (early: RT > 0.5, late: RT < −0.5) (**Fig. 1C**). In contrast, 21.2% and 6.3% of recent enhancers were consistently early and late replicating, respectively, as were 31% and 1.4% of conserved enhancers (**Fig. 1D**). To investigate the emergence of new enhancers, we focused on the averaged replication times across the four germline assays.

The fastest rates of enhancer turnover occurred at late DNA replicating domains. New enhancers were proportionately 1.8 times more common at late than early replicating regions, although the absolute number of enhancers was higher at early replicating domains (**Fig. 1E-G, Fig. S1**). Enhancer turnover was highly correlated with germline replication time (R^2^ = 0.95), although similar trends were observed comparing using somatic developmental replication time (R^2^ = 0.60, **Fig. 1F**, **Fig. S2-3**). It is unlikely the observed trend is due to an ascertainment bias. Beyond the analysis steps taken to ensure the reproducibility of the peak calls (**Methods**), we identify the same relationship if we restrict to mouse-specific enhancer chromatin marks at uniquely mappable coordinates that exhibit sequence conservation across species, thereby excluding potential mappability differences, which could confound the result (**Fig. S4**).

We next examined poised and active enhancers across four mouse tissues (brain, liver, muscle, testis). Recently evolved enhancers were consistently later replicating for both poised and active enhancers across the four mouse tissues (**Fig. 1H**). We found that liver and testis enhancers evolved significantly faster than brain and muscle, with the greatest disparity in enhancer turnover rates between organs at late-replicating regions (*t* = 6.77 and 4.85; p = 1.2×10^−07^ and 3.07×10^−05^ for poised and active enhancers, respectively) (**Fig. 1I**). This result parallels the faster evolution of testis gene expression levels and a slower evolution of brain expression in mammals ^32^. The rapid turnover observed in the testis was also recovered by comparing using somatic replication time (**Fig. S3**).

As transposable elements (TEs) are widespread across the genome and have been widely implicated in the turnover of *cis*-regulatory elements ^5,9^, we next assessed the relationship between TE evolution and replication timing. We calculated the ratio between the numbers of new TE families and the numbers of ancestral TE subfamilies across replication timing regions. Similar to enhancers, new TE subfamilies of TEs were more abundant in late-replicating regions (R^2^ = 0.96, **Fig. S5**). Recently evolved enhancers were also more likely to overlap lineage-specific TEs (**Fig. 1J**). Excluding enhancers overlapping TE (∼52% of enhancers) only slightly reduced the slope between enhancer turnover and replication time (R^2^ = 0.94 and 0.95 for enhancers overlapping and not overlapping TE, respectively) (**Fig. S6)**. Hence, the rate of enhancer turnover is not wholly dependent on TE, but both are strongly correlated to DNA replication time across large chromatin domains ^9,30,33^.

To specifically assess the population of species-specific enhancers that could have emerged from recent copy number duplication events, we grouped human and mouse lineage-specific enhancers separately based on their sequence similarity (**Methods**). While copy number variants arising from recent homologous recombination events or transposition events are expected to share similarity and group, sequences emerging from mutations of ancestral sequences should not. The degree of inter-enhancer similarity depends also on various parameters, including mutation rate, life history, evolutionary pressures, and the evolutionary comparison used for enhancer classification.

Using a significance cut-off of E = 1 × 10^−6^ and a relaxed sequence coverage threshold of greater than 20% of the query sequence to detect homology among recently evolved enhancers, we find the proportions of singleton enhancers are 75.92% and 77.04% for human and mouse enhancers, respectively. Proportions of singletons were similar between human and mouse despite a higher number of expected mutations in mouse. As expected, fewer singletons overlapped repetitive elements, including TE, compared to non-singleton enhancers (Fisher’s exact test, p = 3.56×10^−40^, odds ratio = 0.13 for human enhancers; Fisher’s exact test, p = 6.8 × 10^−171^, odds ratio = 0.63 for mouse enhancers) (**Fig. S7**). Our results are consistent with the idea that most of these elements did not emerge from recent duplication events.

Enhancer gains are more prevalent in regions that already show enhancer marks or chromatin accessibility in other organs ^9,34^. Hence, we repeated our analyses using different thresholds of defining enhancer conservation (conserved enhancer defined as present in at least three or seven other species), which revealed highly consistent results (**Fig. S8**). In summary, although most enhancers are found within early germline replicating domains, species-specific turnover was disproportionately enriched in late replication regions. This was the case for both active and poised enhancers. Late germ line DNA replication time is associated not only with increased numbers of lineage-specific enhancers but also new subfamilies of TEs (not shown). However, most lineage-specific enhancers do not share high degrees of similarities, suggesting the gain of enhancer-associated histone modifications by mechanisms other than duplications.

### Mutations at TF binding sites are linked to enhancer turnover

TF binding sites can be considered the atomic unit of regulatory element function ^35,36^. When mutations occur at TF binding sites, they can disrupt or alter the binding of the TF, potentially leading to changes in enhancer activity. Simulation studies have shown new enhancers can evolve within a relatively short evolutionary time due to the accumulation of mutations creating new TF binding sites^12^.

We hypothesized that the creation or disruption of TF binding sites could change the activity of enhancers, leading to turnover. As TF binding motifs do not fully explain binding, to test this, we used experimental data from TF ChIP-seq to train a deep-learning model to predict binding sites. The model takes a 500 bp DNA sequence and outputs a prediction of TF binding based on sequence alone. We expect a higher predicted frequency of TF binding at new enhancers (i.e., those with recently acquired enhancer histone marks) compared to orthologous but non-enhancer sequences.

Our model architecture was trained on human and mouse ChIP-seq-derived liver-specific TF binding sites, CEBPA, and HNF4A. As above, enhancers were defined based on the enrichment of H3K27ac and the absence of H3K4me3 (**Methods**) ^3^. To optimize the learning of shared functional sequences, the model predicts TF binding using a domain adaptive step to remove sequence biases arising from the species-specific genome backgrounds ^37^. We retrained this model to ensure enhancer regions used for model testing are excluded from the model training process by removing regions harboring human and mouse-specific enhancers. We then tested human and mouse lineage-specific enhancer sequences to assess whether sequence changes could explain enhancer turnover through their impact on TF binding.

The sequence-based model identified a significantly higher number of HNF4A and CEBPA TF binding sites at human enhancers compared to the mouse orthologs without enhancer marks, suggesting that genetic variation between the sequences is associated with the gain or loss of functional TF binding sites (p = 5.78 x 10^−30^; OR = 1.95) (**Fig. 2A, Fig. S9**). Conversely, a similar trend was observed when comparing mouse-specific enhancers to orthologous non-enhancer sequences in human (p = 1.27 x 10^−77^; OR = 3.82) (**Fig. 2A, Fig. S10)**. Enhancer turnover was correlated to sequence changes to the canonical binding motifs (**Fig. 2B)**. Moreover, total proportions of species-specific enhancers with predicted HNF4A and CEBPA binding sites were increased at late replicating regions (**Fig. 2C).** Our findings reveal that mutations altering TF binding can modulate enhancer chromatin states.

**Figure 2.**
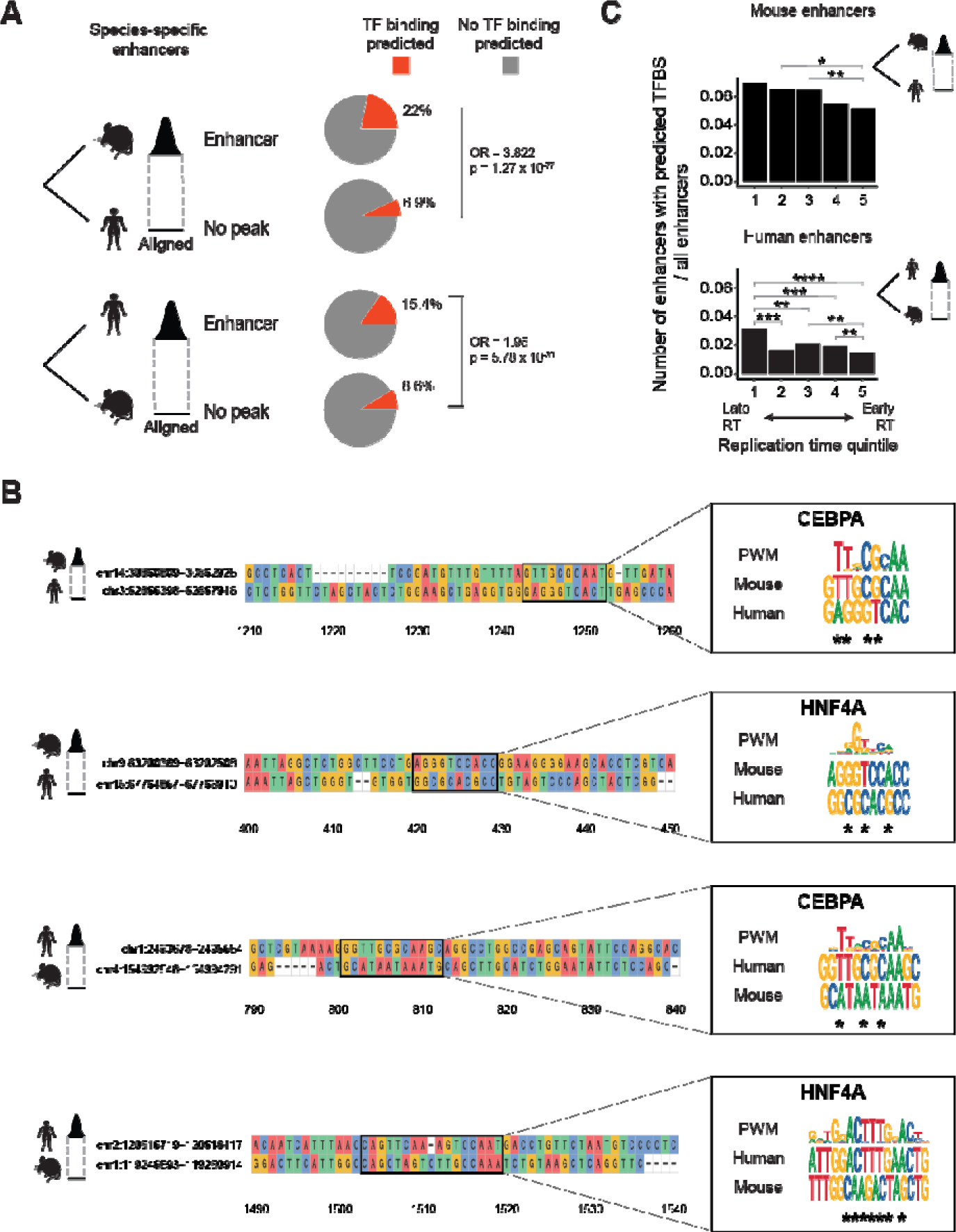
Deep learning model links changes in TF binding sites with enhancer turnover. (**A)** Deep learning domain adaptive model trained with HNF4A and CEBPA binding sites in mouse and human genomes ^37^. Prediction on species-specific enhancers and their aligned non-enhancer sequences in the other species. The pie charts show the percentage of enhancers and matched non-enhancer regions with predicted HNF4A and CEBPA TFBSs with a probability threshold 0.9. Fisher’s exact test odds ratio and p-value are shown for each enhancer vs non-enhancer comparison. (**B)** Examples of species-specific liver candidate enhancers and their sequence alignments to the other species where binding is not predicted in (**A)**. Boxed alignment of a motif identified in the species possessing the enhancer (top sequence) and its alignment to the species without the enhancer (bottom sequence). The motif’s position-weighted matrix (PWM) logo is on the right. The logo is on the negative strand in the last example. **\*** denote changes to PWM in the orthologous sequence without peak; Details on the data processing of this figure is available in **Supplemental Methods**. (**C)** Numbers of mouse-and human-specific enhancers with predicted TFBSs divided by the total number of enhancers across replication time quintiles. The difference in enhancer proportions was tested using Fisher’s exact test between all pairs of DNA replication time quintiles, testing for a higher proportion in the latest quintile (alternative = “greater”). Significance notation: ‘***’ 1 × 10^−4^ < P _≤_ 1 × 10^−3^; ‘****’ P _≤_ 1 × 10^−4^.

### New enhancers are enriched in eQTLs but lack strong signatures of purifying selection

To understand the selective pressures at enhancers, we used human population variation data to calculate a derived allele frequency (DAF) score in 10 bp windows across the genome using whole-genome sequencing of the relatively isolated Icelandic population (deCODE)^38^. DAF odds ratio (OR) measures the ratio between the numbers of rare and common variants. A high odds ratio indicates an excess of rare variants compared to the background, suggesting purifying selection. We plotted DAF for species-specific and conserved liver enhancers by centering each enhancer based on functional motifs to increase the power to detect purifying selection (**Fig. 3A-B**, **Methods**).

**Figure 3.**
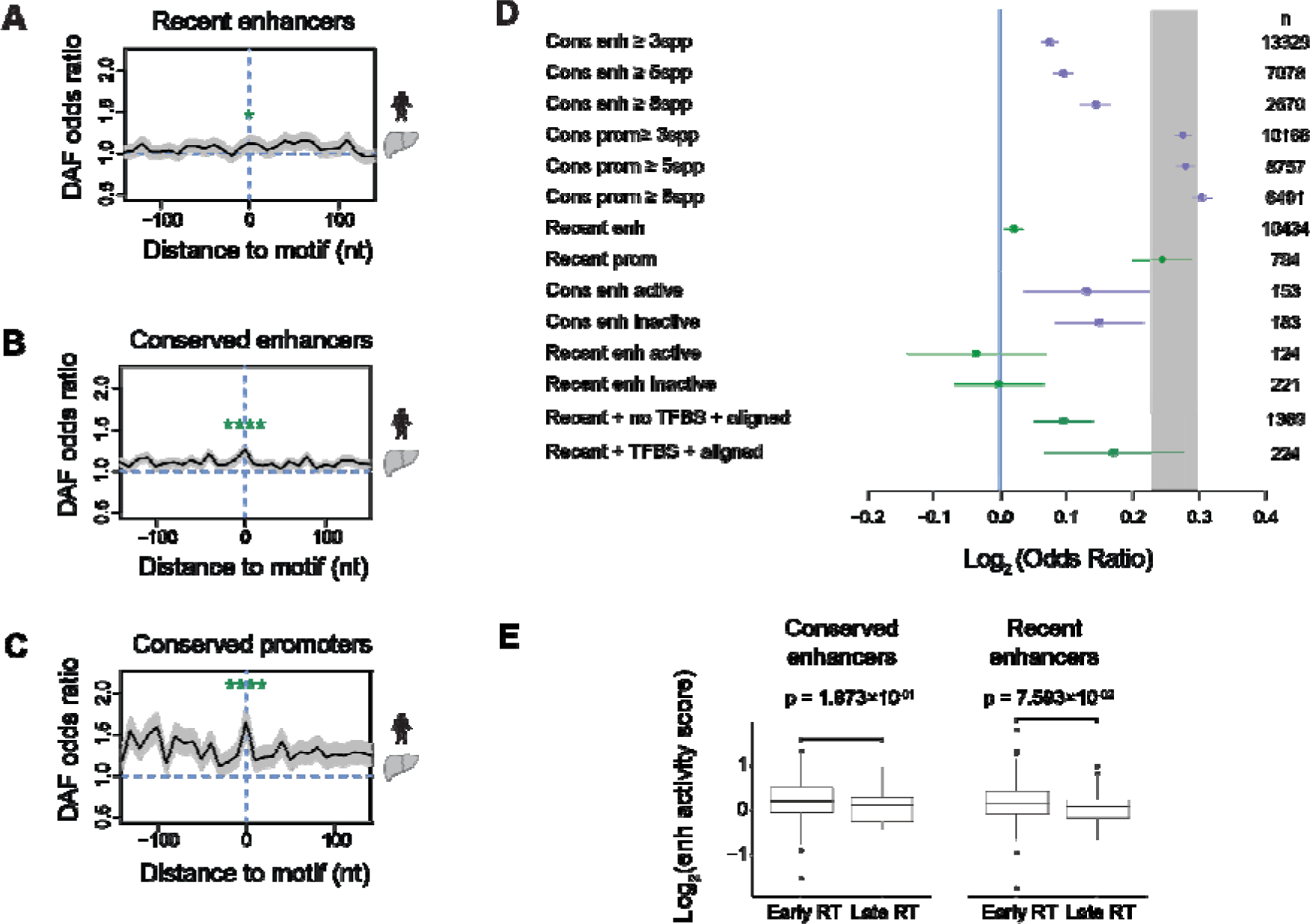
Enhancers do not show strong signatures of purifying selection. Derived Allele Frequency (DAF) odds ratio for recently evolved (**A)** and conserved human liver enhancers (**B)** and conserved promoters (**C)** compared to background genomic regions as a measure of selection pressure. Promoters and enhancers were centered based on the location of liver-specific functional motifs (Fisher’s exact test, significance code ‘*’ P _≤_ 0.05 and ‘****’ P _≤_ 0.0001). (**D)** Log_2_-transformed Odds Ratio of DAF scores for conserved and recent enhancers and promoters. Conservation was defined using multiple thresholds (number of species). Active and inactive enhancers were separated using STARR-seq scores to measure enhancer activity in HepG2 cells ^39^ (**Methods**). DAF Log ORs for human recently evolved enhancers that aligned to the mouse genome where TFBS were detected or not detected using the deep learning model trained for HNF4A and CEBPA in Fig. 2 are shown. Error bars represent the 95% confidence intervals of Fisher’s exact test. Numbers of elements are shown on the right. (**E)** Log_2_ transformed STARR-seq activity of human liver recent and conserved enhancers separated into early (RT > 0.5) and late (RT < −0.5) replicating. Mann-Whitney *U* test p-value is shown in each case.

Our results reveal reduced purifying selection at recent enhancers relative to evolutionarily conserved enhancers and recent promoters. (**Fig. 3C, Fig. S11A-B**). As expected, a progressive increase in DAF OR at enhancers and promoters was observed with increased degrees of species conservation (**Fig. 3D**). Although the DAF scores significantly differed from genome background, this difference may also be due to a higher frequency of common variants rather than a depletion of rare variants (**Fig. S11C-D**). Such differences can be due to demographic and not selective factors. For example, rare variants may not have had as much time to increase frequency and spread through the population, particularly for recently evolved elements.

The low DAF odds ratios suggest many of these newly gained ChIP-seq peaks at late-replicating regions may not be as functional in driving gene expression as their early-replicating counterparts. To delve deeper, we tested whether ChIP-seq peaks at late-replicating regions were as likely to activate transcription as early-replicating regions. Using enhancer activity data from human liver enhancers defined by H3K27ac marks tested in HepG2 cells using STARR-seq ^39^, we compared the normalized activity score between recent and conserved human liver enhancers (recent n = 254, conserved n = 270). We observed slightly lower activity as measured by MPRA at late replication time, although this was not statistically significant (alpha = 0.05) (**Fig. 3D**). We further assessed whether enhancers with MPRA activity showed evidence of increased purifying selection compared to tested enhancers without an appreciable level of enhancer activity (‘inactive’) ^39^. Conserved enhancers with MPRA activity showed a similar DAF odds ratio to conserved enhancers that do not show MPRA activity (**Fig. 3D**). A similar pattern was observed for recently evolved enhancers, where MPRA activity did not distinguish between within species constraint (**Fig. 3D**). The slightly lower measure of activity at late replicating enhancers could reflect a subtle transcriptional function such as weak affinity binding^40^.

To address whether new and conserved enhancers make qualitatively different contributions to transcription, we tested the relative enrichment of 226,768 significant liver *cis*-eQTLs from healthy individuals in the GTEx consortium (GTEx V7) at human-specific and conserved enhancers. Species-specific enhancers harbored significantly more eQTLs than conserved enhancers (Fisher’s exact test, p < 2 × 10^−16^, odds ratio = 1.2), consistent with an increased frequency of eQTLs at recently evolved promoters in the human genome ^41^. Thus, new enhancers harbor more eQTLs than conserved enhancers, suggesting they contribute to regulating gene expression but may be less important for organismal fitness.

### Tissue-specific evolution of enhancers is linked to late DNA replication timing

Late replicating regions with their dynamically regulated heterochromatin and nucleosome formation potential have been linked to tissue-specific gene expression ^24,25,42,43^. Hence, we investigated whether tissue-specific enhancer activity was also associated with late-replicating regions.

Comparing mouse enhancers between the four mouse tissue types, we found tissue-specific enhancers were indeed more likely to be late replicating than enhancers active in more than one tissue (Fisher’s exact test, p = 2.2 × 10^−16^, odds ratio = 0.28, **Fig. 4A**). Late replication time is associated with increased tissue specificity regardless of evolutionary age (**Fig. 4A**). Tissue-specific elements are enriched at late replication time and a faster evolutionary rate than enhancers active in multiple tissues (regression test for difference in slope, p = 7.45 × 10^−6^; **Fig. 4B)**.

**Figure 4.**
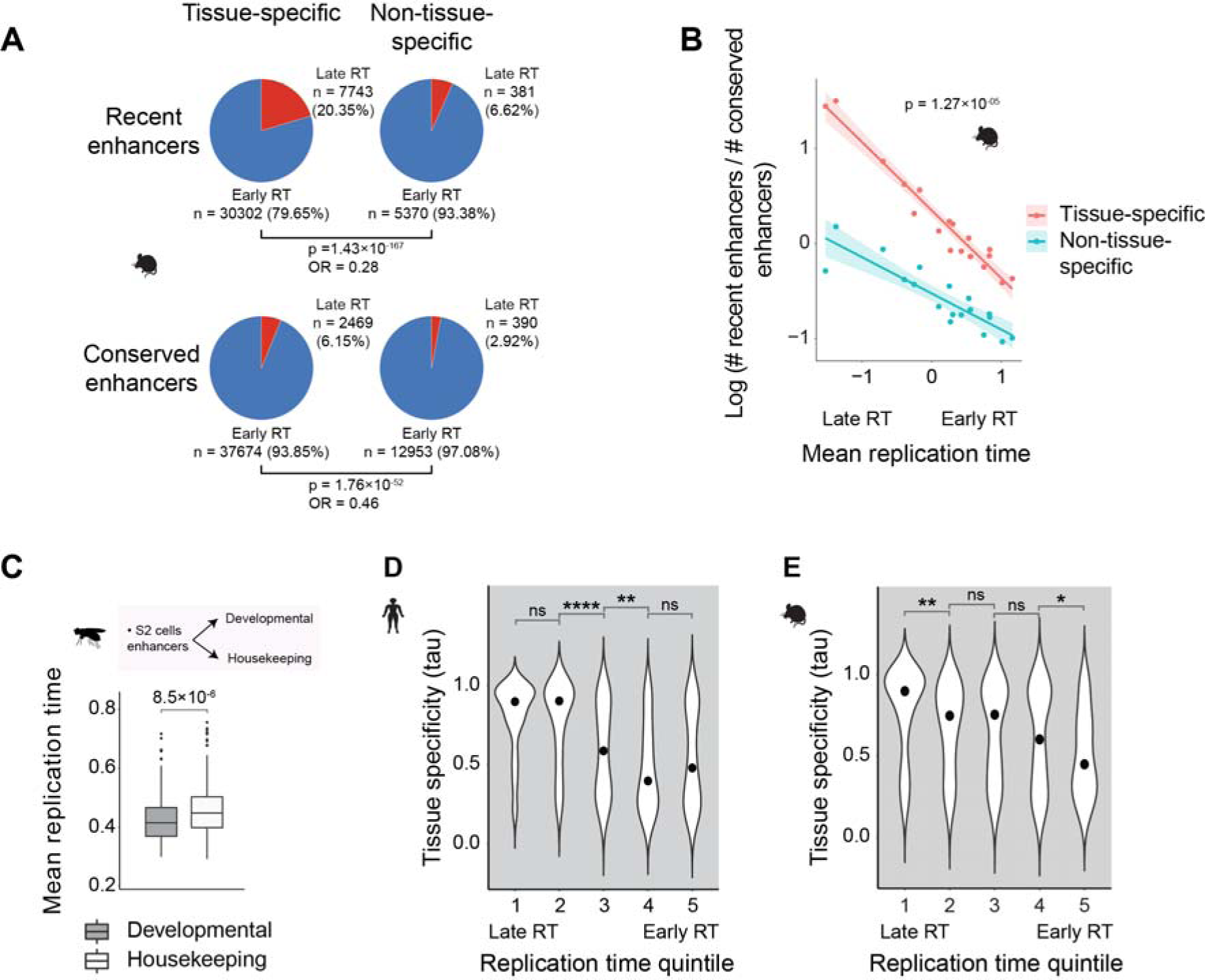
Tissue-specific enhancers are enriched at late replicating regions. (**A)** The proportions of early and late replicating enhancers for tissue-specific and non-tissue-specific mouse enhancers (defined with four tissues: brain, liver, muscle, testis) (Fisher’s exact test, p = 2.2 × 10^−16^, odds ratio = 0.28). (**B)** Mean mouse germline DNA replication time versus enhancer turnover rate, defined as log (number of recent enhancers/number of conserved enhancers), for tissue-specific and non-tissue-specific enhancers (shown in red and blue, respectively) across the 18 DNA replication time clusters shown in Fig. 2C. R^2^ = 0.95 (p-value = 6.43 × 10^−12^) and 0.78 (p-value = 1.19 × 10^−06^) for tissue-specific and non-tissue-specific enhancers, respectively. ANCOVA p-value for the difference in slope is shown. (**C)** Mean replication time of developmental and housekeeping fruit fly enhancers (Mann-Whitney *U*-test, alternative = “greater,” n = 200 enhancers each class). (**D, E)** Violin plots of tissue-specific expression scores (tau values) of human and mouse TFs separated into five quintiles depending on their respective motif enrichments at early versus late replicating enhancers (Mann-Whitney *U*-test, pairwise comparison of later vs. earlier replicating quintile, alternative = “greater,” significance code: ‘ns’ P > 0.05, ‘*’ P _≤_ 0.05, ‘**’ P _≤_ 0.01 and ‘****’ P _≤_ 0.0001). Across the panel, mouse germline replication times are calculated as the mean across PGC and SSC cells, and human replication times are from H9 cells.

Because tissue-specific control of gene expression is critical during development, we hypothesized that enhancers that drive developmental expression may replicate later than those that drive housekeeping expression, which are more likely to be constitutively expressed. Indeed, enhancers associated with developmental promoter activity in *Drosophila* were later replicating than enhancers associated with the housekeeping promoter (**Fig. 4C**; Mann-Whitney *U*-test, p = 8.5 × 10^−6^; **Methods**). This pattern was consistent across different chromosomes independent of the promoters’ endogenous location (**Fig. S12**).

We then examined whether enhancers located at late replicating regions were also associated with binding more tissue-specific TFs. Using an established index of tissue-specificity of gene expression, tau ^44^, we examined tissue-specific TF expression for 477 and 360 genes in humans and mice across 27 and 19 tissues, respectively ^45^. TFs were partitioned into five groups based on the relative enrichment of their motifs at enhancers from early and late replication time. TFs whose motifs were most enriched motifs at late replicating enhancers were significantly more likely to show tissue-specific expression patterns (**Fig. 4D, E**).

### Developmentally associated motifs were enriched at late DNA replication time

In mammals and other warm-blooded vertebrates, DNA replication time is also linked to long regional stretches of compositionally homogeneous DNA with uniform GC base composition ^26,27,46–49^. These are known as GC isochores and are distinct between early and late replicating regions ^48,50,51^. The origin of isochores can be partially explained by mutational biases ^2,26,27^. Late replicating sequences harbor a biased substitution pattern towards A and T nucleotides ^21,52,53^, where the primary contributor is the deamination of methyl-cytosine at CpG sites, resulting in C > T transitions.

Given the nucleotide differences in TF binding sites, we sought to understand how the GC isochore may impact genome-wide TF binding dynamics. GC isochores are closely correlated to replication timing. We confirmed that the loss of G and C nucleotides was greatest at late DNA replication time using base substitutions inferred from the last common ancestor of *Homo* and *Pan* (**Fig. 5A**). We then focused on enhancers and promoters. The general trend of GC content across replication times was similar for cis-regulatory elements and the genomic background, where enhancers with the highest GC content were located at the earlier replicating regions (**Fig. 5B**). Due to CpG islands, promoters possessed the highest GC content levels, but enhancers contained higher GC than genomic background.

**Figure 5.**
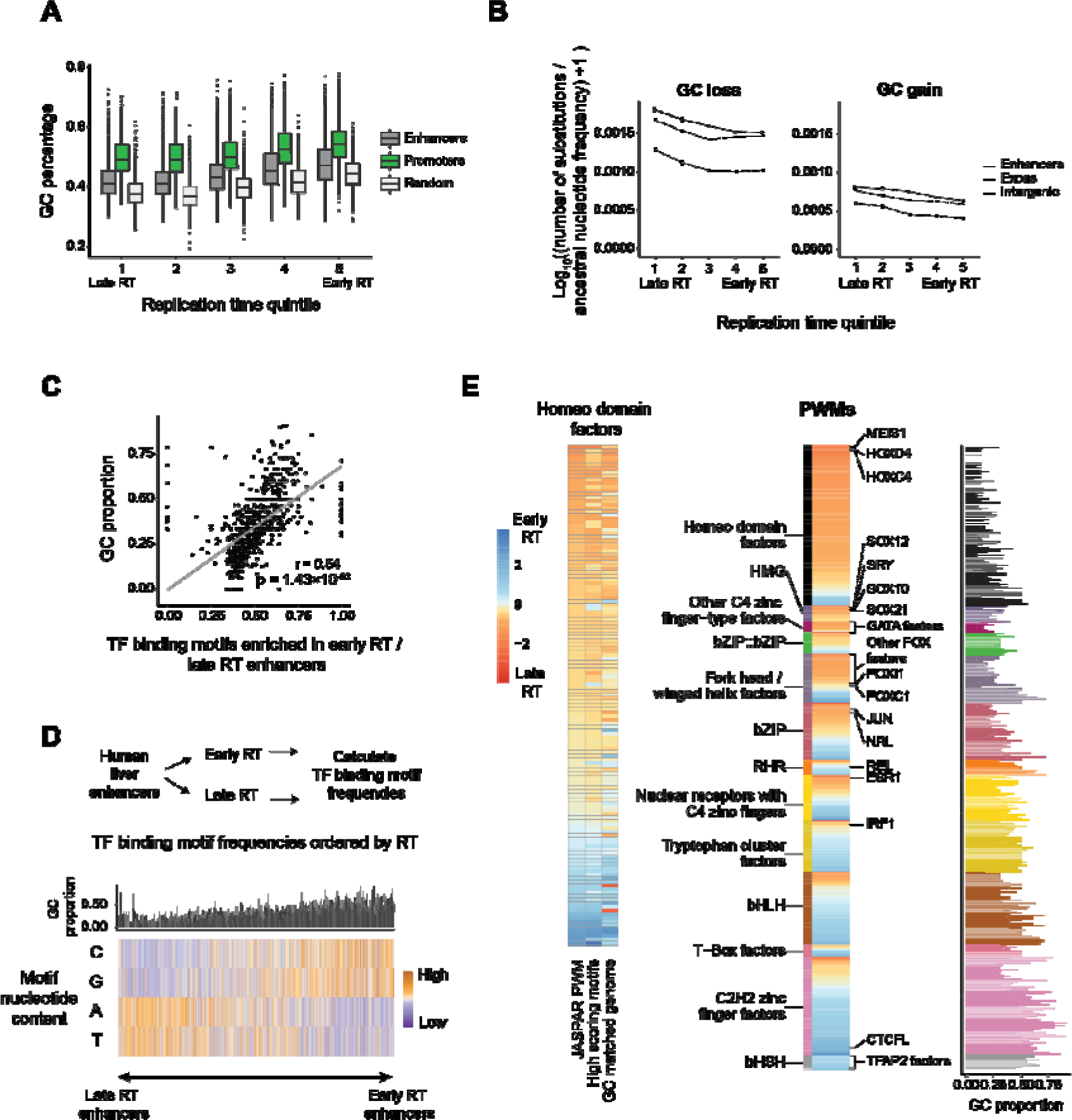
AT-rich motifs are associated with developmental TFs and are overrepresented at late replication time in mammals. (**A)** GC percentage of human liver enhancers and promoters and random genomic regions across replication time quintiles (random regions were sampled from the non-genic areas of the genome, also excluding liver promoters and enhancers) (n = 28175, n = 11520 and n = 5000 enhancers, promoters, and genomic background, respectively). Difference in GC percentage across DNA replication time quintiles was significant for every type of sequence (Kruskal-Wallis’ chi-squared = 2708.6, 688.25, and 902.98 for enhancers, promoters, and random sequences, respectively; p-value < 2.2×10^−16^ for enhancers and p-value = 1.22×10^−147^ and 3.76×10^−194^ for promoters and random sequences, respectively). (**B)** Mean non-CpG substitutions at liver enhancers, exonic, and intergenic regions across H9 replication time quintiles. Substitutions are between humans and the inferred common ancestor of *Homo* and *Pan.* The number of substitutions was adjusted by their ancestral nucleotide frequency, and log_10_ transformed. Error bars represent standard error. (**C)** Scatterplot of the proportion of GC for TF binding motifs (each dot) based on relative enrichment at early versus late replicating human liver enhancers. Motifs enriched within late replicating enhancers are on the left, and those increased at early replicating enhancers are to the right. Pearson correlation r and p-value are shown. (**D)** The bar plot shows the GC proportion of each motif. Heatmap of the GC/AT nucleotide content proportion of TF binding motifs ordered based on their relative enrichment at early versus late replicating human liver enhancers (n = 5538 each replication time). Each column shows a human TF binding motif from the JASPAR database). (**E)** Relative enrichment of TF binding motifs at early versus late replicating liver enhancers grouped by TF class (center heatmap). The GC content of the motifs is shown on the right. Bars are coloured by TF Class. Only TF classes with more than ten TFs are shown. The heatmap on the left shows the relative enrichment of homeo domain factors in early versus late replicating enhancers using JASPAR human motifs (left column) and using only the highest scoring motifs (mid column) (**Methods**). The column on the right shows the relative enrichment of homeo domain factors in early versus late GC%-matched random regions of the genome.

GC isochores corresponded to a profound shift in the counts of different TF binding motifs at enhancers across replication times (**Fig. 5C-D, Fig. S13**). In humans and mice, the most prevalent motifs at late replicating enhancers were AT-rich, while early replicating enhancers were GC-rich (**Fig. 5C-D, Fig. S14**). Homeodomain factor motifs, which act as critical regulators in development, were predominantly enriched in late-replicating enhancers (**Fig. 5E**). A similar trend was observed at promoters (**Fig. S15**). Restricting to the top-scoring motifs resembling the consensus homeodomain TFBSs did not change the observed trend (**Fig. 5E**). To interrogate this further, we also compared TF motif enrichments at regions randomly sampled from the genome (**Fig. 5E**). Motif enrichment was well predicted by replication time, but replication timing did not explain all enriched motifs (e.g., HOXC13, HNF1B, POU4F3) (**Fig. S16**). Some motifs possess a different nucleotide composition than predicted by replication time, suggestive of natural selection.

Because the nucleotide frequencies of motifs at enhancers are an indirect measure of TF binding, we asked whether the observed trends are reflected *in vivo*. Using the DNA binding locations of 71 proteins from ChIP-seq data in human K562 cells, we found TF binding sites were bimodally distributed with respect to replication timing (**Fig. S17**). We fitted Gaussian components using mixture modeling for each protein, focusing on binding sites at later replicating time, which were more variable between TFs than early replicating TFBS (**Fig. S17A-B, Supplemental Table S2**). Our results suggest that DNA replication time impacts the type and frequency of TF binding motifs, thereby influencing the TFs bound at these regions (**Fig. S17C**). However, the trend from *in vivo* binding data is attenuated compared to motifs identified by computational search. The discrepancy could be due to two factors: first, TF binding does not always accurately reflect enhancer locations, and second, TF binding itself relies on more than just the presence of sequence motifs and is influenced by the presence and cooperation of other TFs, as well as the overall arrangement of motifs.

In summary, DNA replication time is associated with not only the tempo/rate of enhancer evolution but also impacts TF motif enrichment.

### Enhancer turnover in cancer is enriched at late DNA replication time

Finally, we asked whether the relationship between DNA replication timing and cis-regulatory element turnover is consistent across evolutionary time scales. Studies have suggested that DNA replication time between healthy and cancer cells is largely stable ^59,60^. Hence, we compared enhancer gains and losses between healthy and cancer cell states across four cancer types across DNA replication time (**Fig. 6A**). We defined a ‘gain’ of enhancers as those characterized in cancer cell lines but not in the non-diseased state. Inversely, enhancers in the healthy cell state but not in cancer were defined as ‘lost.’ Enhancers annotated in both states were termed ‘unchanged’ (**Fig. 6A-B, Supplemental Table S3**).

**Figure 6.**
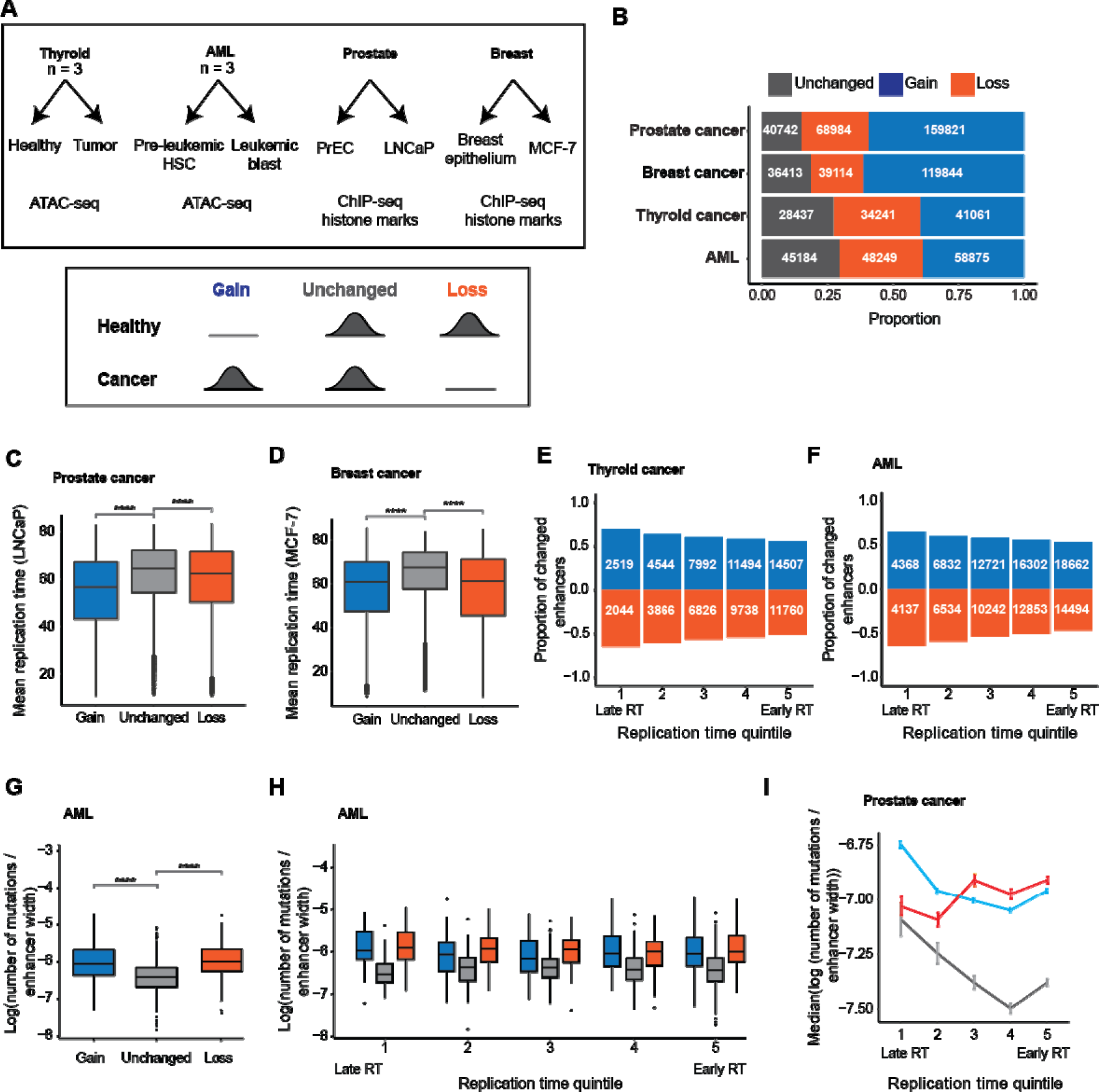
Enhancer turnover is enriched at late replication time in cancer. (**A)** Overview of the enhancer data sets in cancer and matched healthy tissues and cell lines (top). Gained, unchanged, and lost enhancers were defined in each cancer type (bottom). (**B)** Proportions of unchanged, gained, and lost enhancers in each cancer type. (**C-D)** Replication time of gains, unchanged enhancers, and losses in prostate (**C)** and breast (**D)** cancer (Mann-Whitney *U*-test). A similar trend exists in thyroid cancer and AML (**Fig. S13 B and C**). (**E-F)** Proportions of enhancer gains and losses in thyroid cancer (**E)** and AML (**F)** are relative to the number of unchanged enhancers across replication time quintiles (Fisher’s exact test, alternative = “greater,” compared to unchanged enhancers). Proportions of losses were multiplied by (−1). (**G)** Log transformed number of mutations normalized by enhancer width for AML gains, losses, and unchanged enhancers (Mann-Whitney *U*-test). (**H)** Log transformed number of mutations normalized by enhancer width in AML across replication time quintiles (Mann-Whitney *U*-test**). (I)** Median log-transformed number of mutations normalized by enhancer width at prostate cancer gains, unchanged enhancers, and losses across replication time quintiles (error bars represent standard error). Significance notation: ‘ns’ P > 0.05; ‘***’ 1 × 10^−4^ < P _≤_ 1 × 10^−3^; ‘****’ P _≤_ 1 × 10^−4^. Numbers of enhancers and mutations in each cancer type are in **Supplemental Table S3**. Cell type-specific replication timing datasets are used (**Methods**).

We annotated candidate enhancers in healthy breast, prostate, thyroid, and preleukemic cells and their diseased state ^54–58^. In breast and prostate cancer, enhancers were defined by ChIP-seq of histone marks (**Fig. 6A, Fig. S18A, Supplemental Table S3**). In AML and thyroid cancer, enhancers were defined as distal chromatin accessible in patient-matched primary tissues and tumors (thyroid cancer and matched healthy n = 3, pre-leukemic and matched blast cells n = 3). We used DNA replication time information for prostate cancer cell line (LNCaP), healthy prostate epithelial cells (PrEC), and breast cancer cell line (MCF-7) ^59^, with predicted DNA replication time in pre-leukemic and thyroid cells (**Methods**).

Consistent with cross-species results, we found the highest rate of enhancer turnover at late DNA replicating domains compared to enhancers that remained unchanged (**Fig. 6C-F, Fig. S17B-G**). This trend was unaffected by differences in recombination breakpoints ^61^ (**Fig. S19**). Subsequently, we compared cancer variants at gained, lost, and unchanged enhancers in thyroid, AML, and prostate cancer. We used matched tumor and healthy samples from the same individual to calculate somatic mutations due to cancer. Prostate cancer variants were identified from the whole-genome sequencing of the prostate cancer genome cell line, and common population variants were removed ^62^.

Mutation numbers were elevated for enhancers gained or lost compared to unchanged enhancers across the three cancer types for which we had variant data (**Fig. 6G-I, Fig. S17H-N, Supplemental Table S3**). The trend was consistent across all individuals for thyroid cancer and AML (**Fig. S20**). These results illustrate a consistent pattern across different evolutionary scenarios: a higher turnover of cis-regulatory elements at late DNA replication, strongly correlated to an increased mutational burden.

## Discussion

In this study, we demonstrated the significance of genome structure on enhancer evolution. While most enhancers, defined by histone mark occupancy, were identified in early replicating regions, comparative analyses showed that young enhancers were almost twice as likely to replicate later than conserved enhancers. Genetic changes during evolution can create or abolish TF binding sites associated with the emergence of cis-regulatory elements or decommissioning existing elements. The short length of TF binding motifs and their degeneracy allows for the rapid emergence and fixation of TF binding motifs ^11–16^. We found that enhancer turnover is linked to sequence changes that alter TF binding. Remarkably, similar patterns in cis-regulatory evolution were evident in mammalian evolution, spanning millions of years, and in cancer cells, occurring over months or years, suggesting that regulatory evolution is intertwined with the evolution of genome architecture across time scales.

Our definition of species-specific enhancers depended on the other species in the comparison. The closest relatives to humans and mice used in our comparisons were macaque and rats, respectively, with divergence times of ∼29 (human vs. macaque) and ∼12 (mouse vs. rat) million years ago ^63^. Based on the mutation rate and generation time for each species, we should expect ∼15 mutations per kb in humans and ∼120 mutations per kb in mice since the last common ancestor of human/macaque and mouse/rat, respectively (based on mouse mutation rate of 5 x 10^−9^ per base per generation, human mutational rate of 1.28 x 10^−8^, generation time of 0.5 and 25 years ^64,65^). This means the recently evolved mouse enhancers used in the species evolutionary study will show greater genetic variation overall than human-specific enhancers. These mutational differences reflect variation in generation time and effective population size between species and should be considered while interpreting the results.

Our results suggest that evolutionary innovation in gene regulatory modules is more likely to emerge from regulatory elements at later replication domains in a tissue-restricted manner. Disparities in enhancer turnover between organs were also most pronounced in late-replicating regions. Late replicating regions are associated with developmental enhancer activity and are enriched for developmentally relevant TF binding sites related to body patterning (e.g. homeobox). Consistent with this, tissue-specific enhancers are enriched at late replicating regions, where they are associated with tissue-specific TFs.

Our findings also provide a compelling explanation for tissue-specific gene expression differences in mammals linked to GC isochores whose position mirrors replication domains ^59,66^. We showed that the highly organized isochore patterns in mammalian genomes influenced the genome location and frequencies of TF binding sites, with specific types of TFs more likely to be recruited at certain replication timing domains. Specifically, tissue-specific TFs are frequently linked to AT-rich, late replicating binding sites, which may explain the observed enrichment of tissue-specific gene expression in GC-depleted regions ^24,25^.

We speculate that transcriptional changes occurring at late replicating domains may have played a pivotal role in the evolution of the bilaterian body plan and embryonic development of multicellular organisms. Notably, replication timing is dynamic between cell types and varies between germline and somatic cell types ^29,67,68^. Approximately 30% of the human genome switches between replication timing domains across 26 human cell lines ^69^. Therefore, enhancers emerging in germ cells at late replication time could shift to earlier replicating domains in differentiated cell types, where they may have a significant influence on gene activity.

## Methods

### Mammalian enhancer annotation

Unless specified otherwise, all analyses were performed on the human and mouse genome assemblies hg19 and mm10. R v4.0.0 ^70^ was used. Species-specific ChIP-seq datasets used are E-MTAB-2633 and E-MTAB-7127 ^3,9^. To summarize, the candidate enhancer identification strategy reads were aligned using BWA v.0.5.9/0.7.12, and peaks were called using MACS v.1.4.2/2.1.1 using total DNA input control with p < 1 x 10^−5^ threshold. Consensus peaks that overlapped two or three biological replicates by a minimum of 50% length were used. Enhancers were defined as those regions that overlapped an H3K27ac or H3K4me1 enriched region but not a H3K4me3 enriched region.

Conserved human enhancers (n = 13329) were defined as liver enhancers in at least two of 18 other mammalian species (Rhesus macaque, green monkey, common marmoset, mouse, rat, Guinea pig, rabbit, Northern tree shrew, dolphin, sei whale, Sowerby’s beaked whale, cow, pig, dog, cat, ferret, opossum and Tasmanian devil) ^3^. Recently evolved (i.e., human-specific) enhancers (n = 10434) were defined as human cis-regulatory elements without a histone mark indicative of enhancer activity in another species at aligned regions (∼85%) or did not align to the genomes of other species (∼15%). **Supplemental Tables S4 and S5** show the mean enhancer width per dataset (human and mouse) and the number of human enhancers aligned to other species’ genomes, respectively.

The alignment of mouse recent enhancers to other species’ genomes was determined based on liftOver mapping with option -minMatch = 0.6. We used chain files for the assemblies RheMac10, CalJac4, Rn6, OryCun2, SusScr11, CanFam3, FelCat9, EquCab3, and MonDom5.

To check the overlap of conserved and species-specific with chromatin accessible regions, we used ATAC-seq data for human liver and DNAse-seq data for mouse brain, liver, and muscle (**Supplemental Table S1**). The minimum overlap of an enhancer with ATAC-seq or DNAse-seq peaks was 30% of the enhancer base pairs in all cases. The alternative hypothesis tested was a higher overlap of conserved enhancers with accessible regions using the option alternative = “greater” in Fisher’s exact test. Conserved enhancers were conserved in at least two other species.

### Replication time data

Repli-seq data was generated by treating cells with 5-bromo-2’-deoxyuridine (BrdU), a thymidine analog, to label newly synthesized DNA. Subsequently, cells are fixed and FACS-sorted based on their DNA content into early S phase and late S phase cell populations. The DNA from these cells is then amplified and mapped to the reference genome. To quantify the timing of DNA replication, the ratio of normalized read coverage between the early and late fractions is calculated ^71^. Higher values in this ratio represent early DNA replication; low values indicate late replication. The replication time of every enhancer was calculated by averaging the replication time of the regions they overlap. Early and late replication elements were denoted as mean times > 0.5 and < −0.5, respectively. Difference in the probability of recent enhancers between the early and late replication time was calculated as follows:

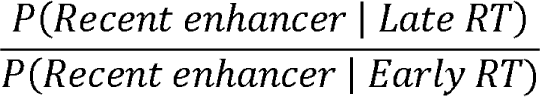

Z-score transformed replication timing data was obtained from human ESC H9 ^72^, mouse primordial germ cells (PGC), and spermatogonial stem cells (SSC) (Acc: GSE109804)^30^, and 22 mouse cell lines differentiated from ES cells ^29^. Mean mouse germline replication time was calculated across PGC (n = 2) and SSP cell lines (n = 2). Mean somatic replication time was calculated across all 22 mouse cell lines. For cancer analyses, DNA replication time information for prostate cancer cell line (LNCaP), healthy prostate epithelial cells (PrEC), and breast cancer cell line (MCF-7) ^59^, together with predicted pre-leukemic and thyroid DNA replication time data using ATAC-seq information was used (see below). All genomic regions with available replication time data were included in downstream analyses.

### STARR-seq data for HepG2

We used STARR-seq data of human liver enhancers defined by ChIP-seq and tested on the HepG2 cell line ^39^. After removing negative controls, we separated the tiles (n = 6735) to active and inactive groups using the published threshold (log_2_ score > 1) and overlapped to our enhancers.

### Estimation of DNA replication time using ATAC-seq

Where relevant replication timing data was unavailable, ATAC-seq data was used to infer replication time using Replicon v0.9 ^73^. ATAC-seq signal was normalized to a mean of 0 and unit variance. Replicon was run with default options on every chromosome (excluding scaffolds). The predicted replication time values were multiplied by −1 to match the direction of Repli-Seq RT values. The mean ATAC-seq signal across pre-leukemic samples was used to predict replication time.

### Clustering of mouse replication time data

Replication time data from 22 early embryonic mouse cell lines differentiated from mouse embryonic stem cells were transferred to mm10 coordinates (USCS liftOver tool) and overlapped with the replication time regions from PGC (n = 2 cell lines) and SSP cell lines (n = 2 cell lines) ^29,30^. Mean replication time was calculated for every 200 kb region across the mouse genome (n = 8966 replication time bins across all cell types). Mean replication time values were centered and scaled using the function “scale” ^70^. The function “Mclust” from the R package mclust was used to estimate the best number of *k*-means clusters of 200 kb replication time bins (G = 1:k.max, modelNames = mclust.options(“emModelNames”), where k.max is 20) (mclust version 5.4.6) ^74^. The best number of clusters was selected based on its Bayesian Information Criterion (BIC) (k = 18, BIC = −37291.04). Replication time bins clusters were obtained with the function kmeans (18, iter.max = 20). Cell types were clustered using hierarchical clustering with k = 9 (function hclust from base R, method = “complete”, distance = “euclidean”).

### Tissue specificity score

Tau (r) scores of tissue specificity were calculated with the following formula:

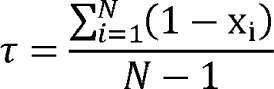

Where N represents the number of tissues, x_l_ represents the expression profile of one tissue divided by the maximum expression value across tissues ^44^. This analysis used the previously described mouse tissue data from the brain, liver, muscle, and testis.

### Model to test for tissue-specific differences

We used a linear model to test for differences in tissue-type-specific evolutionary rates. Using the formula logFC_enh ∼ mean_RT + tissue_pair + tissue_pair:mean_RT, where logFC_enh represents the log (number of recent enhancers/number of conserved enhancers) values, tissue_pair is the tissue pair codified as binary (liver and testis = 1, brain and muscle = 0) and mean_RT is the mean germline replication time. The interaction effect was tested using the t-test statistic.

### Transposable elements

Transposable elements from RepBase (v27.04 for mouse and human)^75^ were annotated using RepeatMasker (v4.0.6 for mouse, v4.1.0 for human) using the sensitive search setting for mouse (‘-s’) ^76^. Species-specific elements, as classified by RepBase, were termed ‘recent,’ and the remaining TEs termed ‘ancestral.’

### Developmental enhancer analysis

Summit coordinates of *Drosophila* enhancers determined using STARR-seq on the S2 cell line were downloaded (Acc: GSE57876)^77^. Housekeeping and developmental promoter were of *RpS12* and *even-skipped* TF, respectively. We defined housekeeping and developmental enhancers as the most highly ranked 200 enhancers for each promoter based on their STARR-seq score. Fly DNA replication time profiles for S-phase in S2-DRSC cells were used (Acc: GSE41350)^78^. Enhancer summits were extended by 250 bp upstream and downstream, and each enhancer’s mean replication time was calculated.

### Cross-species deep learning model

We used the model architecture from Cochran et al. (2022) ^37^ and retrained their model to ensure our test data was not used in the training process and to focus the model on learning differences at orthologous regions that show species-specific histone marks indicative of differences in enhancer activity between human and mouse genomes.

A domain adaptive neural network architecture was used to remove background sequence biases between human and mouse genomes at TF binding sites ^37^. Input data was generated by splitting the mouse (mm10) and human (hg38) genomes into 500 bp windows, with 50 bp offset. After excluding all regions containing human- and mouse-specific enhancers and their orthologous region in the other species, we trained the model described in Cochran et al. ^37^. Liver human and mouse HNF4a and CEBPA ChIP-seq peak data from ^2^ were remapped to hg38 or mm10, respectively. We trained two sets of models for every TF: humans as the source species and mice as the source species. For each species, peaks were converted to binary labels for each window in the genome: “bound” (1) if any peak center fell within the window, “unbound” (0) otherwise. We constructed balanced datasets for training using all bound regions and an equal number of randomly unbound sampled (without replacement). Sequence data was one hot encoded. Human and mouse genome sequences were used for model training, excluding Chr 1 and Chr 2. Genome windows from Chr 2 were used for testing. Genome windows from Chr 1 were used for validation. For each TF and species, models were trained for 15 epochs to reduce bias (**Fig. S21)**. Final models were selected based on maximal auPRCs. The test data set comprised species-specific enhancers centered on the middle 500 bp of each element. For predictions using the model, we used a probability >= 0.9. USCS ‘liftOver’ with minMatch = 0.6 was used for genome assembly remapping. We selected models that maximized the auPRC. We evaluated the performance of the models using test datasets (**Fig. S22**). We used the models to predict TF binding in species-specific enhancers centered on the middle 500 bp of each element.

### Natural selection analysis

Human genome variation data was retrieved from the deCODE whole-genome sequencing study of the Icelandic population ^38^. Derived Allele Frequency (DAF scores) of every segregating SNP was calculated, and alleles were defined as either rare ( < 1.5% population frequency) or common ( > 5% frequency) as previously described ^41^. The number of rare and common alleles in 10 bp windows were centered with respect to the locations of functional liver-specific TF binding motifs from the database funMotifs; 75 unique funMotifs were counted ^79^. These counts were normalized for the average rates with 2–4 kb upstream and downstream flanking regions. Confidence intervals were obtained by performing 100 bootstrap replicates of sampling the motif locations with replacement. Odds ratios of rare against common alleles between enhancers (and promoters) and size-matched background genomic regions selected randomly were calculated in 10 bp windows. Odds ratio confidence intervals and p-values were obtained using Fisher’s exact test. Only autosomes were considered.

### Copy number analyses

Homology was assessed using blastn with the option -max_target_seqs N (blast+/2.11.0)^80^; this option was used to retrieve the maximum number of hits for every enhancer; N represents the number of enhancers in every dataset, 10434 and 80904 for human and mouse, respectively. Hits were filtered by E-value < 1 × 10^−6^ and query coverage > 20. We defined singleton enhancers as enhancers without significant similarity to other enhancers.

### Motif frequency analysis

Motif enrichment in human and mouse enhancers and human promoters used the function annotatePeaks.pl from HOMER (option -size given) with human motifs from the JASPAR 2020 database (n = 810). The reference genome annotation was provided through the option -gtf ^81^. To calculate the nucleotide composition of JASPAR motifs, a nucleotide was assigned to a position of the PWM matrix if its frequency was higher than 0.5. Otherwise, an ‘N’ is set to that position. The proportion of every nucleotide is calculated with respect to the length of the motif (number of bases).

Motif replication time was calculated as the relative enrichment of a motif early against late replicating enhancers. For each motif, we used the formula log_2_ ((present_early / absent_early) / (present_late / absent_late)), where present_early is the number of early replicating enhancers (RT > 0.5) with non-zero motif instances of a given motif and absent_early is the number of enhancers with zero motif instances. Similiarly, present_late and absent_late represent the number of late replicating enhancers (RT < −0.5) with non-zero and zero motif instances, respectively. This measure reflects the relative abundance of the motif between early versus late replicating enhancers. The nucleotide composition of motifs was calculated based on the motif consensus sequence. High scoring homeodomain motifs were defined as the motifs in the top quintile of all human homeodomain motifs according to their score from HOMER annotatePeaks.pl.

We identified genomic regions with matched GC content using the function genNullSeqs from the R package “gkmSVM” (version 0.83.0). GC%, sequence length and repeat content were matched with 2% tolerance (repeat_match_tol = 0.02, GC_match_tol = 0.02, and length_match_tol = 0.02, batchsize = 5000, nMaxTrials = 50).

### Gaussian mixture models

We collected Chip-seq data for the binding sites of 71 transcription factors. Hg19 coordinates were used. We overlapped all the TF binding sites with H9 ESC DNA replication time and calculated each TF binding site’s mean DNA replication time. Afterward, we built a Gaussian mixture model for every TF using the function normalmixEM with k = 2 to get two components (R package mixtools version 1.2.0) ^82^. The function returns mu, sigma, and lambda values for each component. Mu represents the mean DNA replication time; sigma denotes the standard deviation; lambda indicates the final mixing proportions (i.e., the contribution of each component to the final mixture distribution).

### Cancer datasets

ChIP-seq data from the prostate cancer cell line, LNCaP, was used to annotate enhancers (Acc: GSE73783) ^57^. For healthy prostate epithelial cells (PrEC), enhancers were defined using chromHMM (Acc: GSE57498) ^58^. We used ChIP-seq data of histone marks in the breast cancer cell line, MCF-7, to annotate enhancers (Acc: GSE96352, GSE86714) and healthy epithelial breast cells (patients epithelium samples) (Acc: GSE139697, GSE139733). LNCaP, MCF-7, and breast epithelium enhancers were regions enriched in H3K27ac or H3K4me1, excluding proximal regions (± 1kb from TSS).

### Cancer ATAC-seq pre-processing and peak calling

We used matched ATAC-seq from cancer and healthy thyroid samples from three individuals (Acc: GSE162515; C1, C7, C8) ^55^, and ATAC-seq files for the matched pre-leukemic and blast cells from three individuals (Acc: GSE74912; SU484, SU501, SU654) ^54^. ATAC-seq fastq files for the matched cancer and healthy thyroid samples from three randomly chosen individuals were downloaded (Acc: GSE162515; C1, C7, C8) ^55^. Adapter sequences were identified and removed using BBDuk (ktrim = r k = 23 mink = 11 hdist = 1 tpe tbo, http://jgi.doe.gov/data-and-tools/bb-tools/). Trimmed reads for each sequencing run were mapped to genome assembly hg19 with bowtie2 v.2.3.5.1 in paired-end mode^83^. Discordant and poor-quality reads were removed (-f2 -q30 -b), and the output was sorted with samtools v.1.10 ^84^. The obtained .bam files were merged by sample (MergeSamFiles) with duplicates removed (MarkDuplicates, http://broadinstitute.github.io/picard/), resulting in three tumor and three healthy libraries. ATAC-seq fastq files for the matched pre-leukemic and blast cells from three randomly chosen individuals with AML (Acc: GSE74912; SU484, SU501, SU654) ^54^ were processed similarly. For each sample from each cancer type, peaks were called using MACS3 ^85^ (-g hs -f BAMPE -B and default q-value cutoff of 0.05), and a union set of peaks was defined.

### Cancer variant calling

We restricted our mutational analysis to single nucleotide polymorphisms (SNPs). Prior to variant calling in thyroid cancer and AML enhancers, we corrected for systematic bias and other sequencing artifacts. Base quality scores of ATAC-seq reads were recalibrated with BaseRecalibrator and ApplyBQSR (GATK v4.2.5.0 ^86^) based on the known variants in 1000 Genomes and Database for Genomic variants (--known-sites Mills_and_1000G_gold_standard.indels.b37.sites.vcf --known-sites Homo_sapiens_assembly19.known_indels_20120518.vcf --known-sites dbsnp_138.b37.vcf.gz). To distinguish somatic mutations, in addition to using the public variant databases, we generated a custom database from the matched healthy samples to filter out the patient-unique germline variants. This panel of healthy (pon) was developed by calling variants on the healthy samples in Mutect2 ^87^ (with option --max-mnp-distance 0), restricting the analysis to the open chromatin regions identified by MACS3. Using the Genome Aggregation Database (gnomAD) as a public germline mutations source, the variants were compiled into a single pon vcf file. After variant calling in cancer samples, we applied FilterMutectCalls to flag candidates likely to suffer from alignment, strand or orientation bias, and germline mutations, as identified in the general population and by pon. From the obtained filtered vcf files, only variants marked by PASS in the FILTER field were considered for further analyses.

For prostate cancer, variants called from the whole genome sequence of LNCaP ^62^ were mapped from hg38 to hg19 using picard LiftoverVcf (v2.26.10, http://broadinstitute.github.io/picard/). Germline genetic variations found in the population were removed using three datasets: HapMap, 1000 genomes phase 3, and National Heart Lung and Exome Sequencing Project data ^62^. We removed indels using bcftools view (option “--types snps”) (v1.9) ^88^. Where variants with multiple alleles existed, one was selected at random. 48,161 putative somatic mutations were identified across 37,482 enhancers in prostate cancer. Hence, an average of 1.46 mutations was observed per enhancer (0.08% of total sequence length).

## Data availability

Code is available here https://github.com/ewonglab/enhancer_turnover. Datasets are deposited to Zenodo doi:10.5281/zenodo.10494781

## Acknowledgments

We thank M.Roller and P.Flicek for assistance with metadata access, J.Wong, C.Du, S.Clark, and S.Mahony for helpful discussions, Q.Wang for assistance with data processing, and C.Liang for assistance with figure making. PCP is supported by a UNSW International Postgraduate Scholarship. VP is supported by an Australian Government Research Training Stipend Scholarship. ESW is supported by an NHMRC Investigator Grant (GNT2009309), ARC Discovery Project (DP200100250), and a Snow Fellowship.

## Supplemental Methods

### Data processing for constructing Figure 2B

We highlighted specific sequence segments to visualize the motif positions identified by FIMO within the candidate enhancer sequences. For this visualization, we leveraged the R package ggmsa (version 1.3.4) (Zhou et al. 2022). Of note, the segments of putative non-enhancer sequences illustrated in the figure did not yield any instances of mouse or human CEBPA or HNF4A motifs.

The complete sequence of mouse and human candidate enhancers was mapped to either the human or mouse genomes using the UCSC liftOver tool with a minimum ratio of bases that remap (-minMatch) of 0.6. We utilized the mm10 genome assembly for mouse and hg38 for human. To refine the dataset, we identified the mouse and human candidate enhancers overlapping CEBPA or HNF4A binding sites as identified by ChIP-Seq data specific to each species (Schmidt et al. 2010).

We focused on the candidate enhancers overlapping with either CEBPA or HNF4A binding sites for subsequent analysis. We used the FIMO tool to identify CEBPA and HNF4A motif instances in the candidate enhancers overlapping CEBPA or HNF4A binding sites, respectively (Grant et al. 2011). In this analysis, we employed mouse CisBP-2.0 motifs for the mouse candidate enhancers and human CisBP-2.0 motifs for the human candidate enhancers (Weirauch et al. 2014). FIMO was used with default parameters, using a p-value threshold of <= 1 × 10^−04^.

To prioritize candidate enhancers, we ranked them based on their CisBP-2.0 motif scores. Specifically, we selected the candidate enhancer with the highest motif score for each transcription factor and species, provided it surpassed the prediction threshold in the domain adaptive model (predicted probability ≥ 0.9).

The full sequences of the chosen mouse and human candidate enhancers, along with their corresponding mappings to the human or mouse genome, were aligned using Clustal Omega (version 1.2.4) via the EMBL-EBI tool with default parameters (Madeira et al. 2022). The clustalW output format was converted to FASTA format using the EMBOSS Seqret tool (Madeira et al. 2022).

---

**Figure S1.**
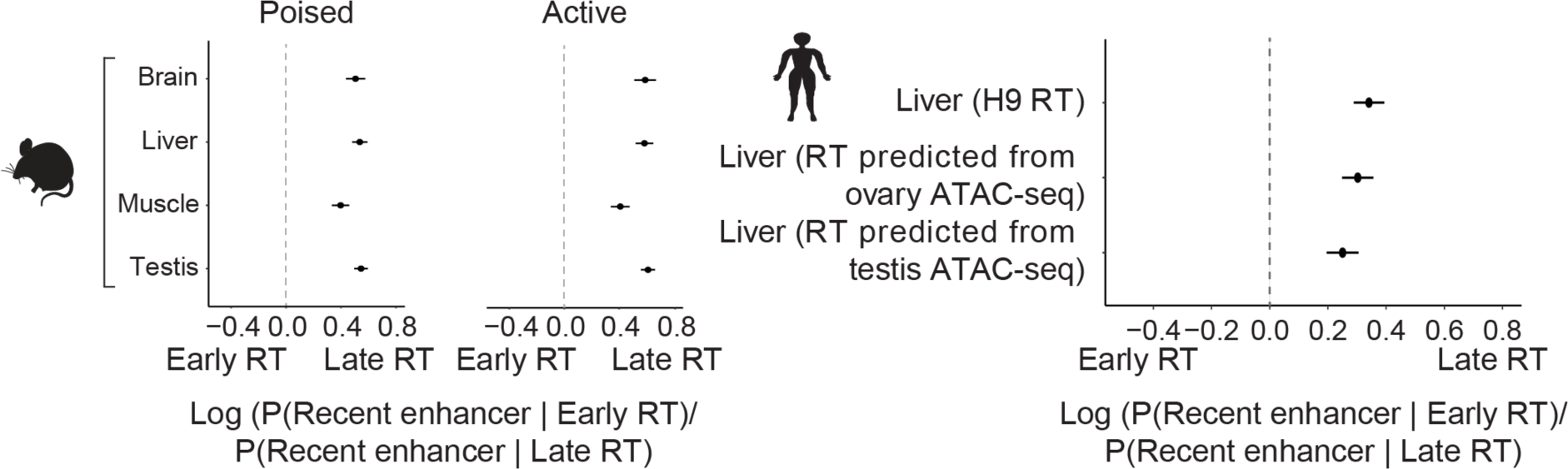
Log odds ratio of the conditional probability of recent liver enhancers dependent on replication time. P(Recent enhancer | Early RT) / P(Recent enhancer | Late RT) values for mouse tissue-specific enhancers separated into poised and active (left panel). Dots represent the mean conditional probability, and error bars are standard errors. Human germline DNA replication time was predicted from ovary and testis ATAC-seq.

**Figure S2.**
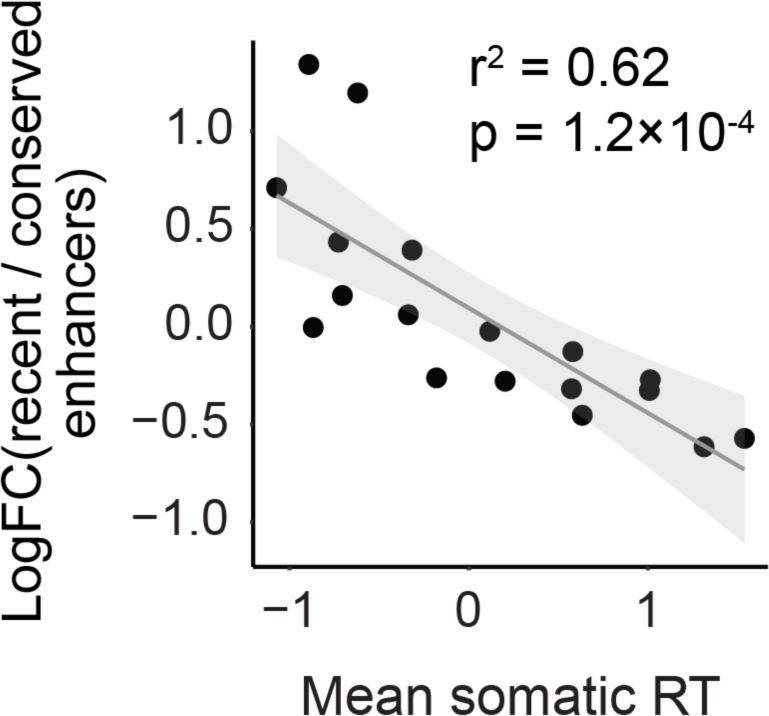
Scatterplot of mouse enhancers turnover and somatic replication time. Scatterplot of mean somatic replication time across the 18 clusters shown in (**Fig. 1C**). R^2^ and p-value are indicated. Mean somatic replication time is calculated across 22 cell lines (**Methods**).

**Figure S3.**
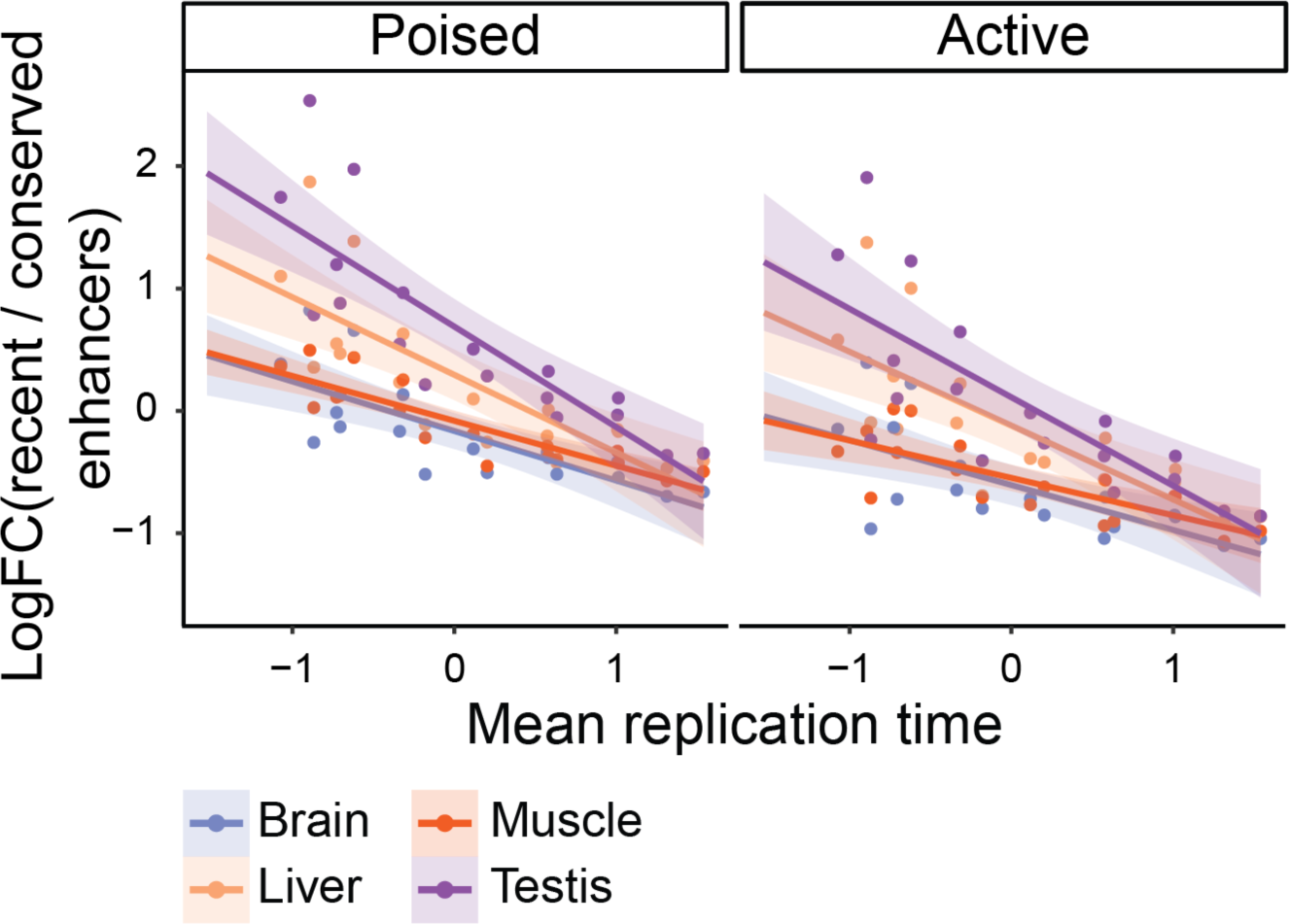
Relationship between mouse enhancer turnover and somatic replication time. Scatterplot showing somatic cell lines mean somatic replication time (x-axis) and enhancer turnover (log FC of recent enhancers against conserved enhancers) by tissue and enhancer type (poised or active) (lm for the difference in slopes between liver and testis against brain and muscle, formula = logFC.enh ∼ mean_rt + tissue_pair + tissue.pair:mean_rt, where logFC.enh is the value of log ( number of recent / number of conserved enhancers), tissue_pair is either “liver_testis” or “brain_muscle” codified as binary, 1 and 0, respectively, and mean_rt is the mean of developmental cell types’ replication time, *t* = −2.23 and *t* = −1.93 for poised and active enhancers, respectively, p = 3.28×10^−02^ and p = 6.31×10^−02^, for poised and active enhancers). Mean somatic replication time is calculated across 22 cell lines (**Methods**).

**Figure S4.**
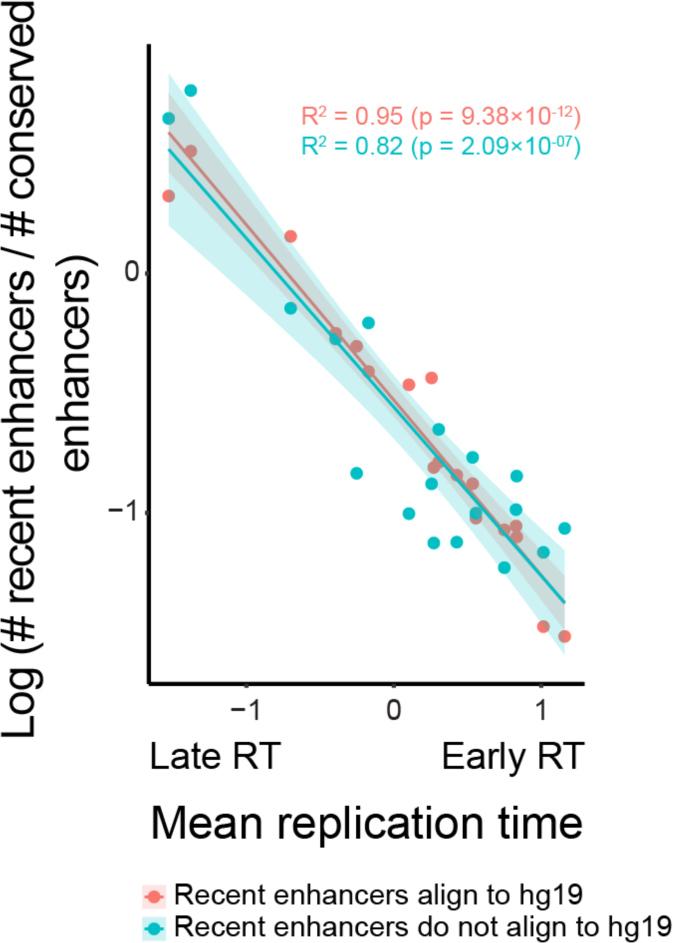
Mean DNA replication time versus enhancer turnover rate for mouse recent enhancers that align and do not align to the human genome. The enhancer turnover rate was calculated using mouse recent enhancers that align and do not align to the human genome (hg19) (liftOver -minMatch = 0.6) for each cluster shown in Fig. 2C. Mean replication time represents mean germ line DNA replication time. R^2^ and p-value are displayed for each group.

**Figure S5.**
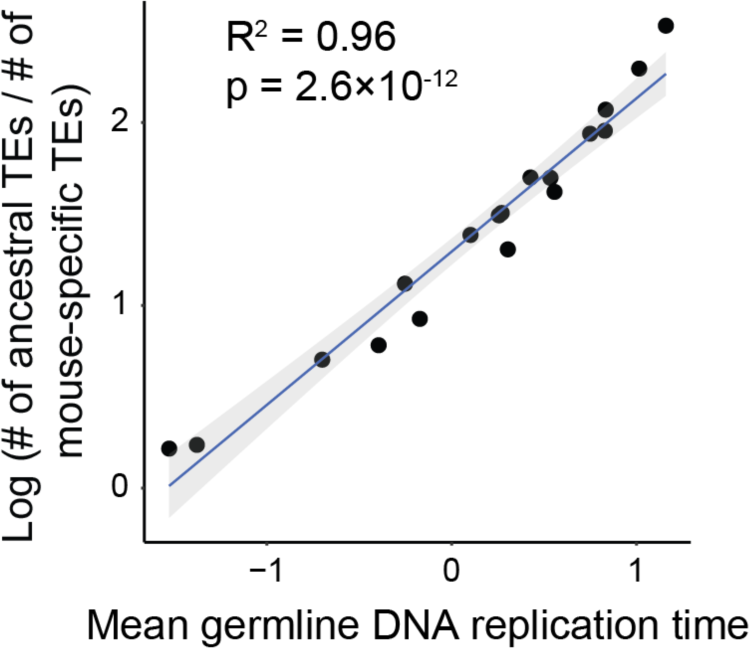
Correlation between mean mouse germ line replication time and TE turnover rate. TE turnover rate and mean germ line DNA replication time are shown for the 18 clusters shown in Fig. 1C. TE turnover rate was defined as log (number of ancestral TEs / number of mouse-specific TEs). R^2^ and p-value are shown.

**Figure S6.**
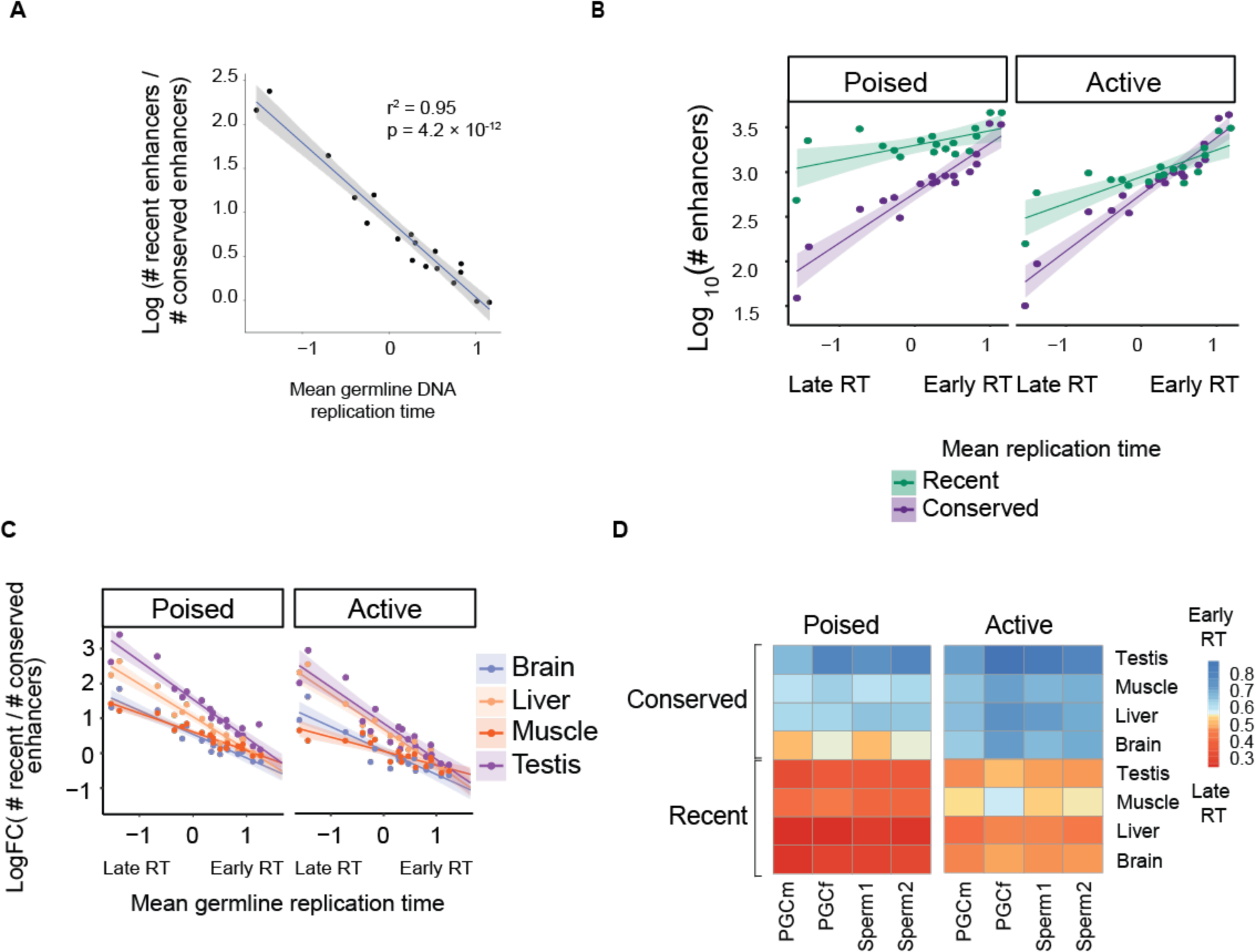
Increasing the number of species used in the definition of evolutionary conserved peaks produced similar outcomes. Evolutionarily conserved enhancers are defined by cross-species conservation among five species (including mouse) (**A)** Scatterplot of mean germline replication time (PGC + SSP) across the 18 clusters shown in Fig. 1C. R^2^ and p-value are indicated. (**B)** Scatterplot of germline mean DNA replication time (PGC + SSP) and log_10_-transformed numbers of recent and conserved enhancers. Each data point represents a cluster as defined in Fig. 1C. (**C)** Scatterplots of germline mean DNA replication time (PGC + SSP) and enhancer turnover by tissue and enhancer type. Each data instance corresponds to a cluster in Fig. 1C. (**D)** Heatmaps of mean PGC and SSC DNA replication time of poised and active mouse enhancers separated by tissue and type.

**Figure S6.**
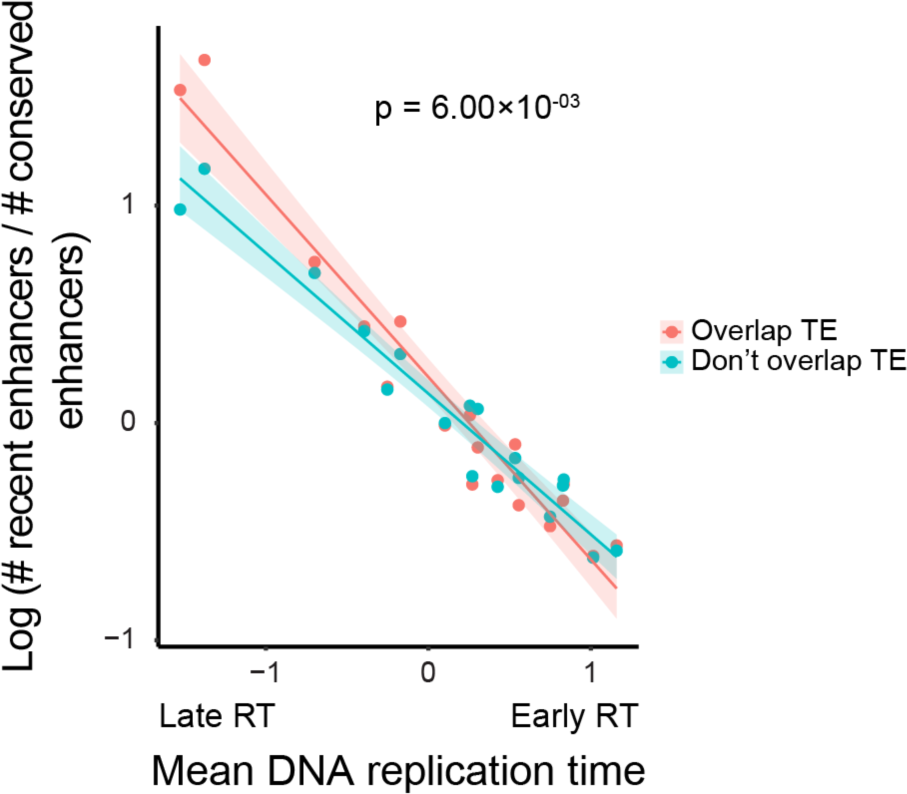
Enhancer turnover rate versus mean DNA replication time for mouse enhancers that overlap and do not overlap TE. Mean germ line DNA replication time against mean enhancer turnover rate (defined as log (number of recent enhancers / number of conserved enhancers)) across 22 DNA replication time clusters as defined in Fig. 1C. Mean enhancer turnover rates and DNA replication time values were separated for enhancers overlapping TE and not overlapping TE (shown in red and blue, respectively). The difference in slope between the two groups was not significant (ANCOVA, p-value shown in the figure). R^2^ = 0.94 (p-value = 2.59×10^−11^) and 0.95 (p-value = 1.13×10^−11^) for enhancers overlapping and not overlapping TE, respectively.

**Figure S7.**
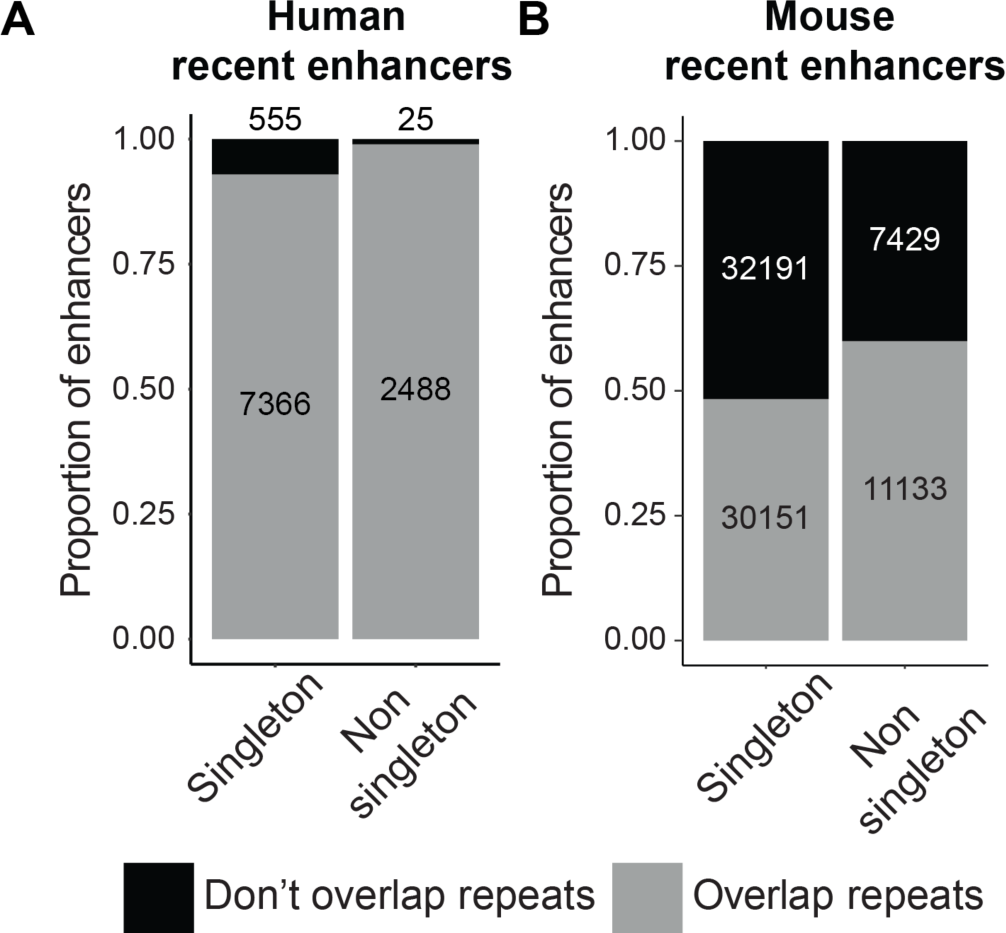
Singleton recent enhancers are less likely to overlap repetitive elements (A) Proportion of human recently evolved enhancers overlapping repetitive elements (Fisher’s exact test, p = 3.56×10^−40^, odds ratio = 0.13). Enhancers are divided into singleton and non-singleton based on cluster analysis (**Methods**). (**B)** Same as in (**A)** for mouse recent enhancers (Fisher’s exact test, p = 6.8×10^−171^, odds ratio = 0.63).

**Figure S8.**
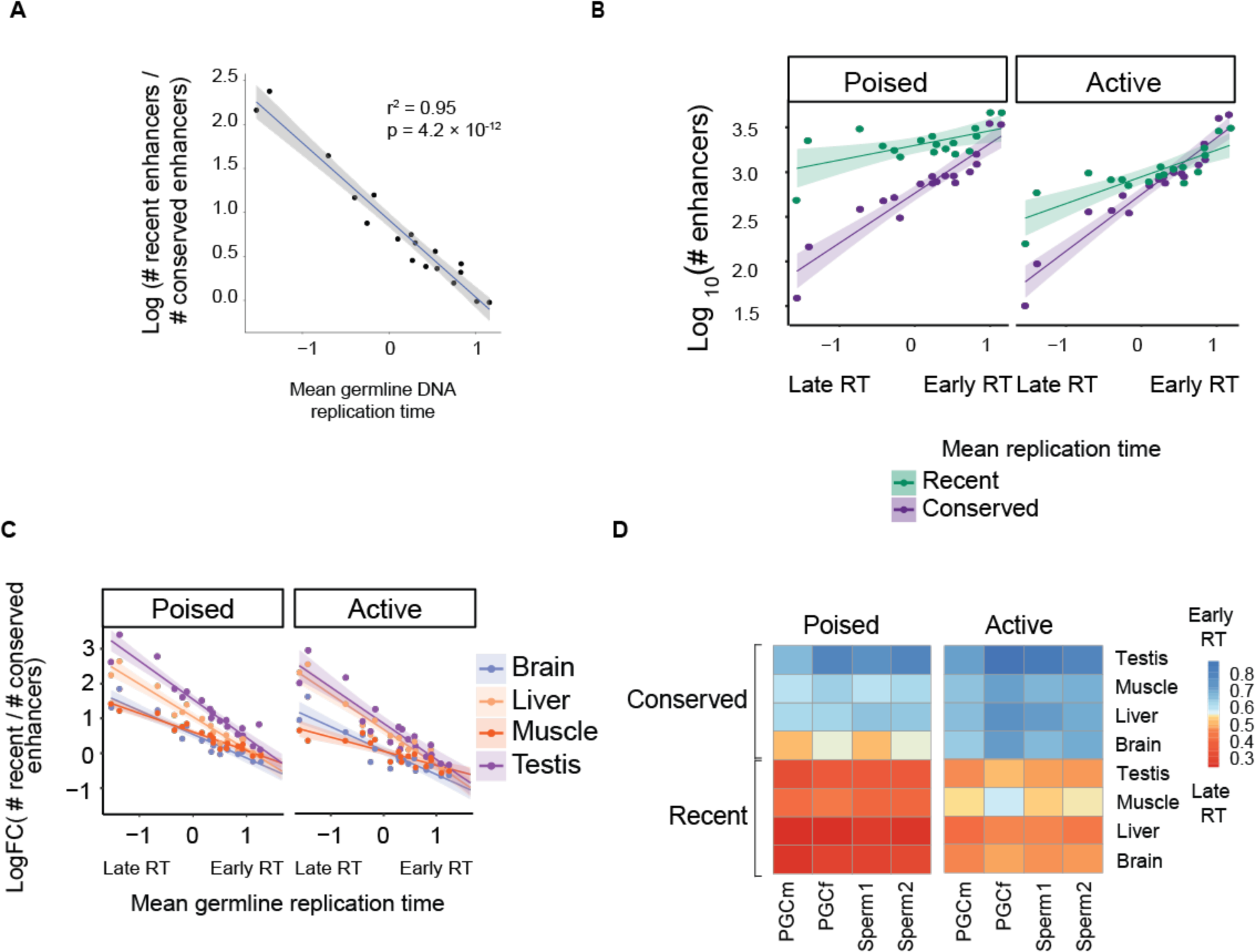
Consistent results with increased stringency in the definition of conserved peaks. Evolutionarily conserved enhancers are defined by cross-species conservation among five species (including mouse) (n = 48580)**. (A)** Scatterplot of mean germline replication time (PGC + SSP) across the 18 clusters shown in Fig. 1C. R^2^ and p-value are indicated. (**B)** Scatterplot of germline mean DNA replication time (PGC + SSP) and log_10_-transformed numbers of recent and conserved enhancers. Each data point represents a cluster as defined in Fig. 1C. (**C)** Scatterplots of germline mean DNA replication time (PGC + SSP) and enhancer turnover by tissue and enhancer type. Each data instance corresponds to a cluster in Fig. 1C. (**D)** Heatmaps of mean PGC and SSC DNA replication time of poised and active mouse enhancers separated by tissue and type.

**Figure S9.**
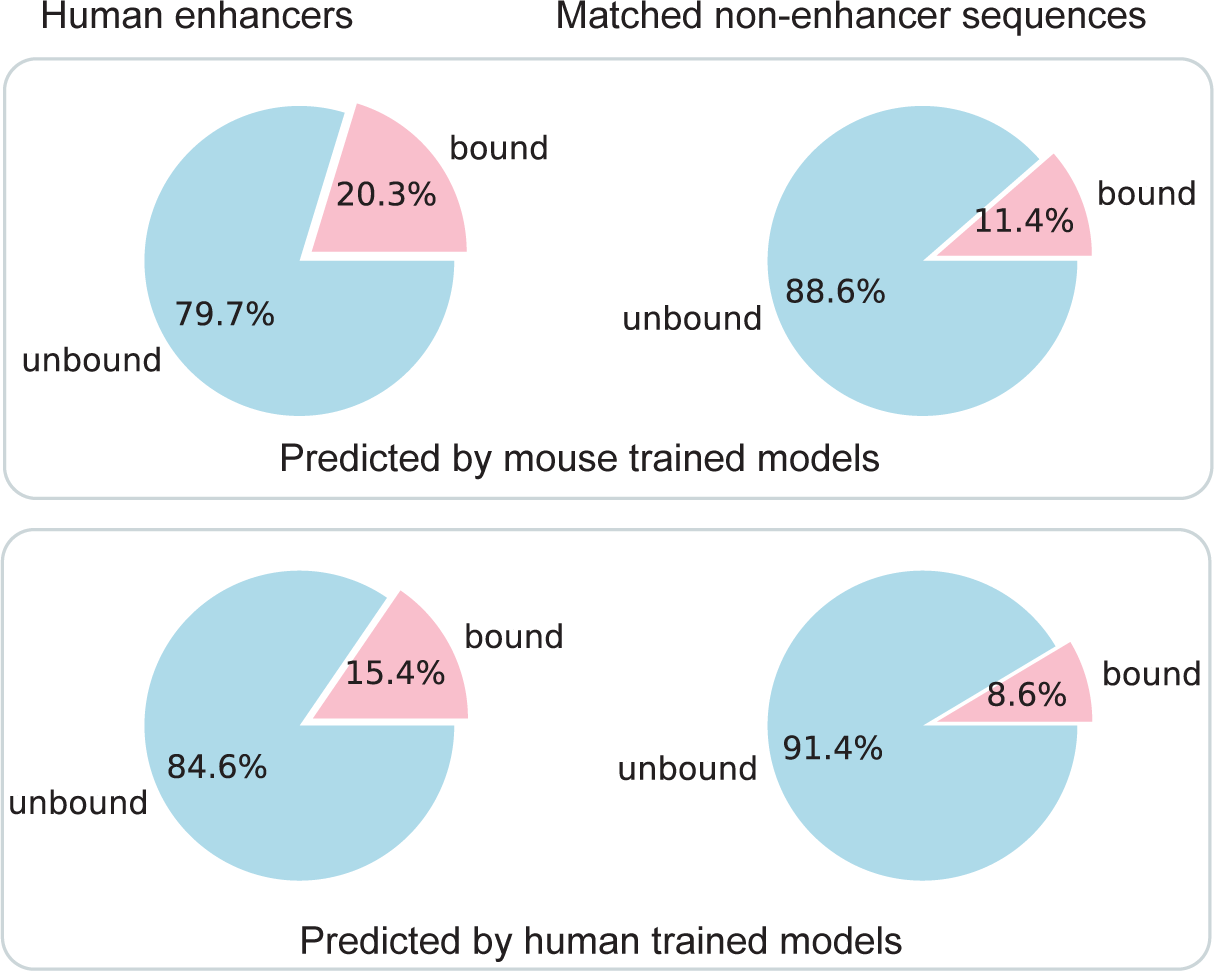
TF binding prediction of human enhancers and orthologous non-enhancer sequences in mouse. The pink sections represent the proportions of human enhancers or non-functional regions predicted to be bound by CEBPA or HNF4A, while the blue sections represent regions not predicted to be bound by any of the TFs. Predictions from models trained on mouse data are displayed on the top, while the bottom row shows predictions from models trained on human data.

**Figure S10.**
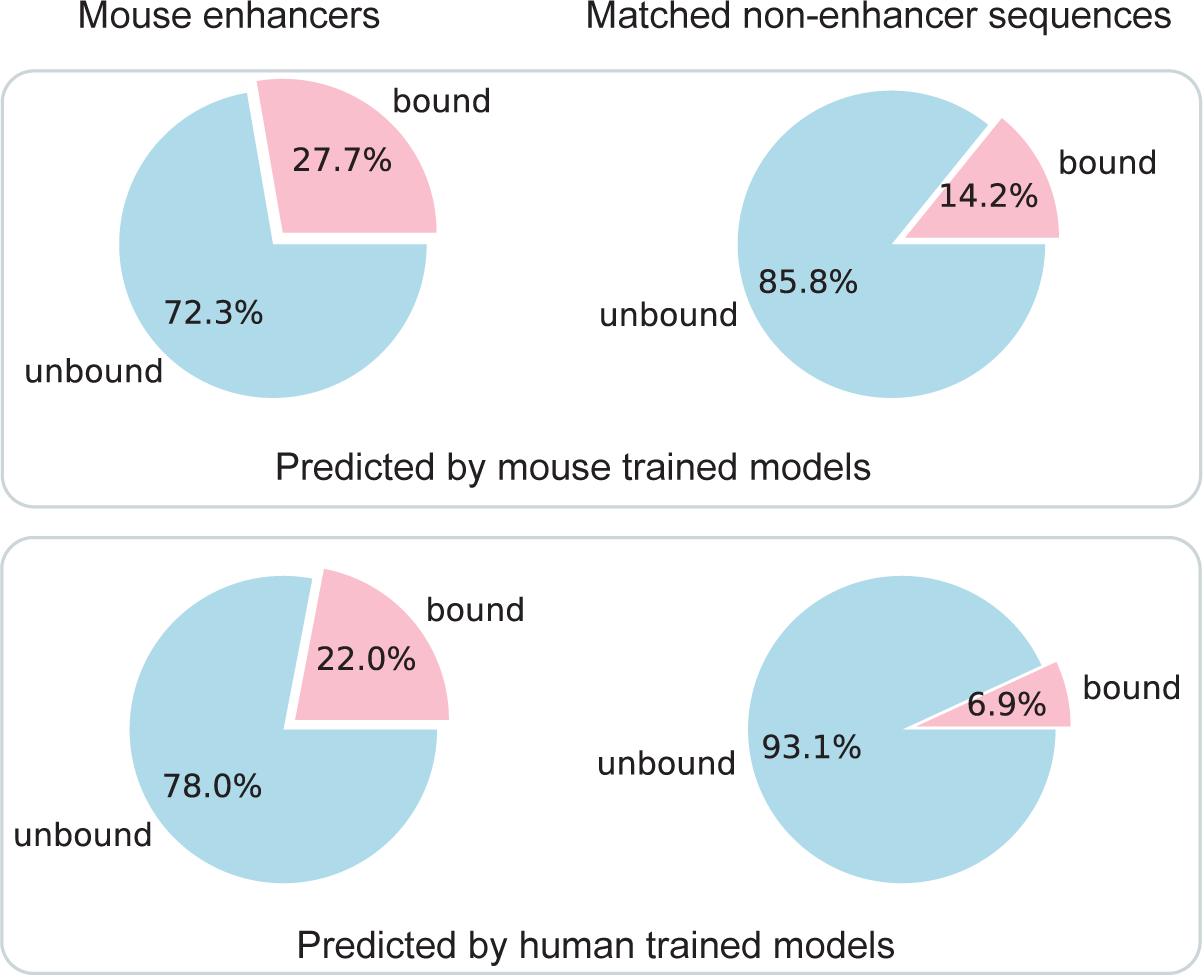
TF binding prediction of mouse enhancers and orthologous non-enhancer sequences in human. The pink sections represent the proportions of mouse enhancers or non-functional regions predicted to be bound by CEBPA or HNF4A, while the blue sections represent regions not predicted to be bound by any of the TFs. Predictions from models trained on mouse data are displayed on the top, while the bottom row shows predictions from models trained on human data.

**Figure S11.**
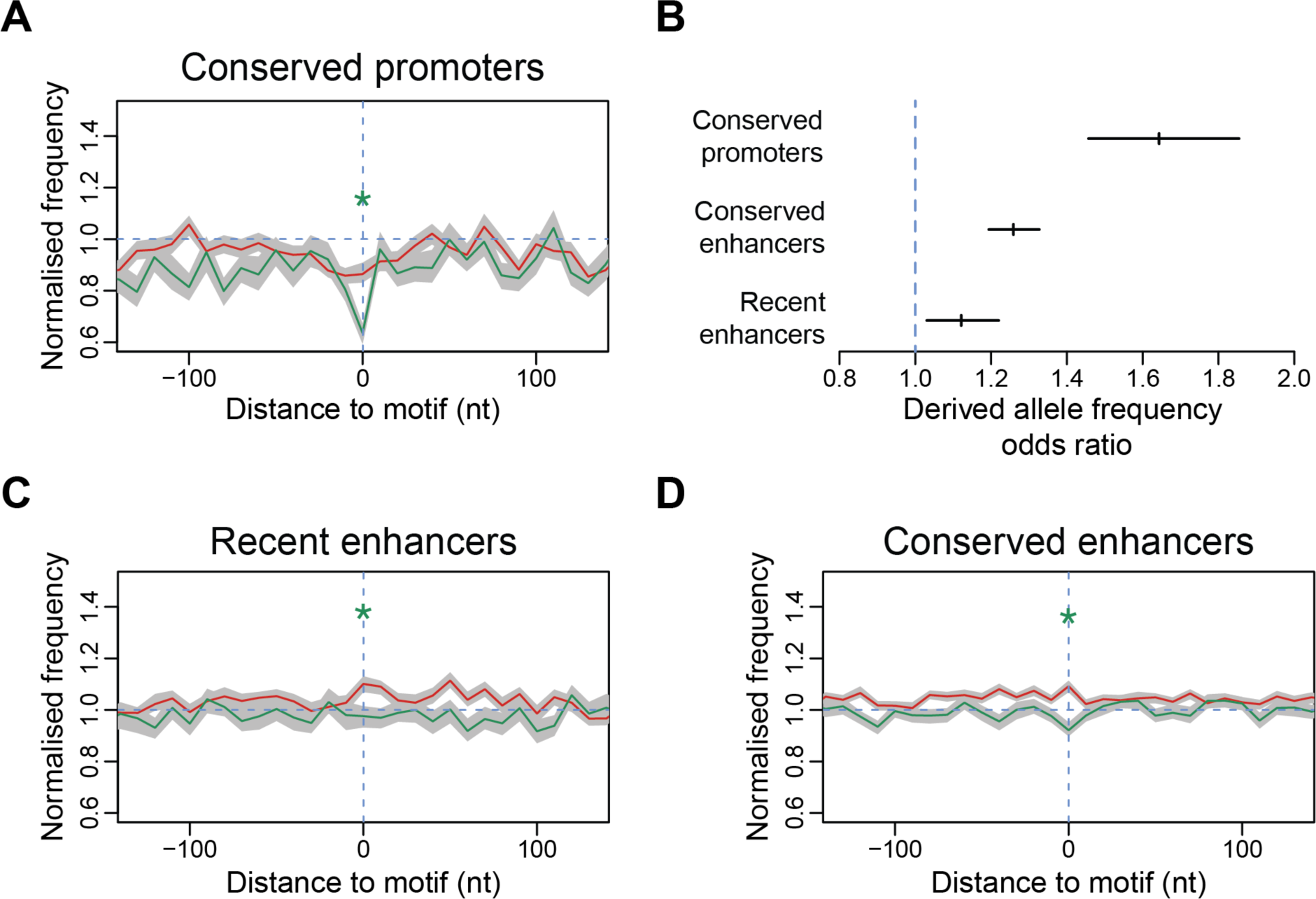
Frequency of rare and common variants at human liver promoters and enhancers (A) Frequency of rare (<1.5% population frequency, red) and common (>5% frequency, green) SNPs at conserved human liver promoters. Promoters were centred based on the positions of functional liver motifs (10bp windows). Allele frequencies were normalised by the average frequencies within 2-4 kb upstream and downstream flanking regions for each category. Shaded areas represent 95% confidence interval obtained by sampling the data with replacement (“*” indicates the absence of overlap of rare and common alleles’ confidence intervals). (**B)** Odds Ratio of Derived allele frequency (DAF) of liver promoters and enhancers. Vertical bars represent DAF odds ratio at recent and conserved enhancers and conserved promoters centred on functional liver motifs (10bp windows) compared to a similar number of windows selected at random from the genome (relative to Fig.2 **F-H**). Error bars represent 95% confidence interval obtained from a Fisher’s exact test. (**C-D)** Similar to (**A)**, the frequency of rare (red) and common (green) allele frequencies at recent (**C)** and conserved (**D)** liver enhancers.

**Figure S12.**
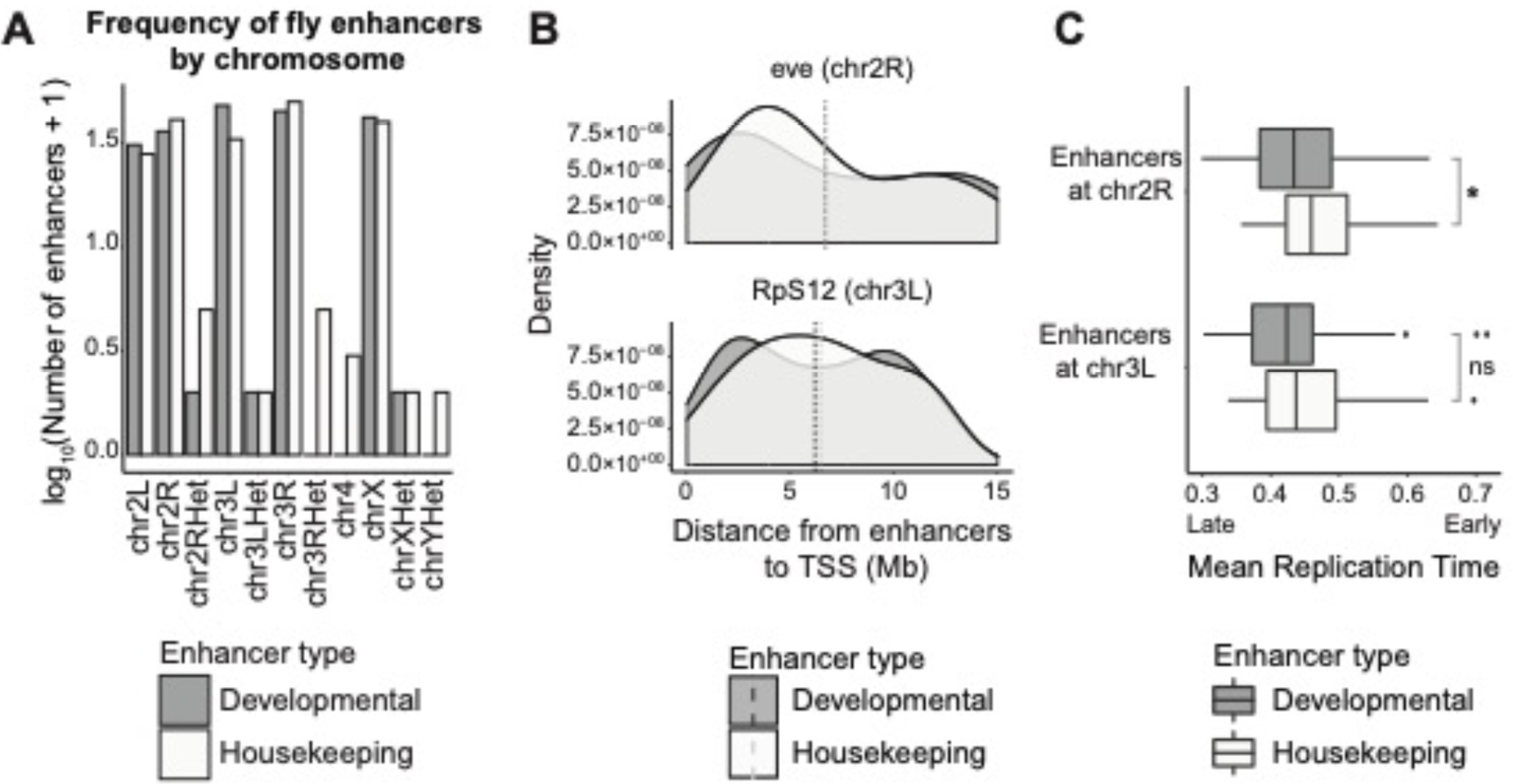
Distribution of fruit fly developmental and housekeeping enhancers. (**A)** Frequency of developmental and housekeeping fly enhancers across the *Drosophila* genome (dm3). (**B)** Distance of developmental and housekeeping enhancers to the Transcription Start Sites (TSSs) of *eve* and *RpS12*, whose promoters were used as developmental and housekeeping promoters, respectively (n = 35 and 40 developmental and housekeeping enhancers at chr2R, n = 47 and 32 developmental and housekeeping enhancers at chr3L). (**C)** Mean replication time of enhancers shown in (**B) (**Mann-Whitney *U*-test, developmental vs housekeeping enhancers, significance code ‘ns’ P > 0.05 and ‘*’ P ≤ 0.05).

**Figure S13.**
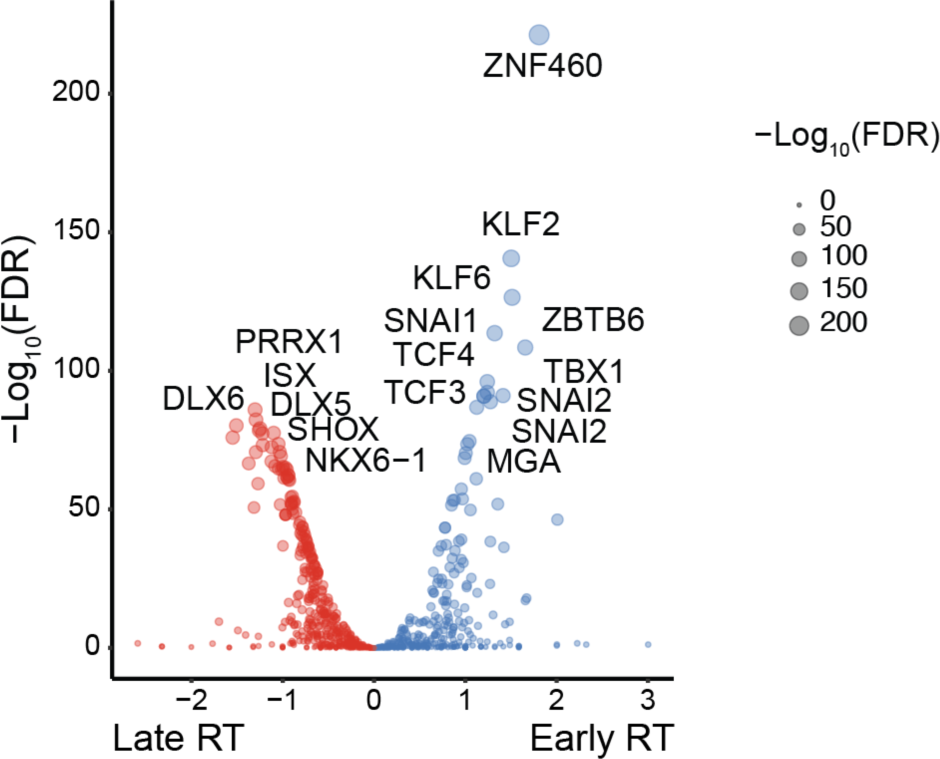
Transcription factor binding sites enriched in early versus late replicating human liver enhancers. Enriched JASPAR motifs between early and late replicating liver enhancers. X-axis shows the relative enrichment for each motif at early versus late replicating enhancers; y-axis represent -log_10_ (Fisher’s exact test, FDR).

**Figure S14.**
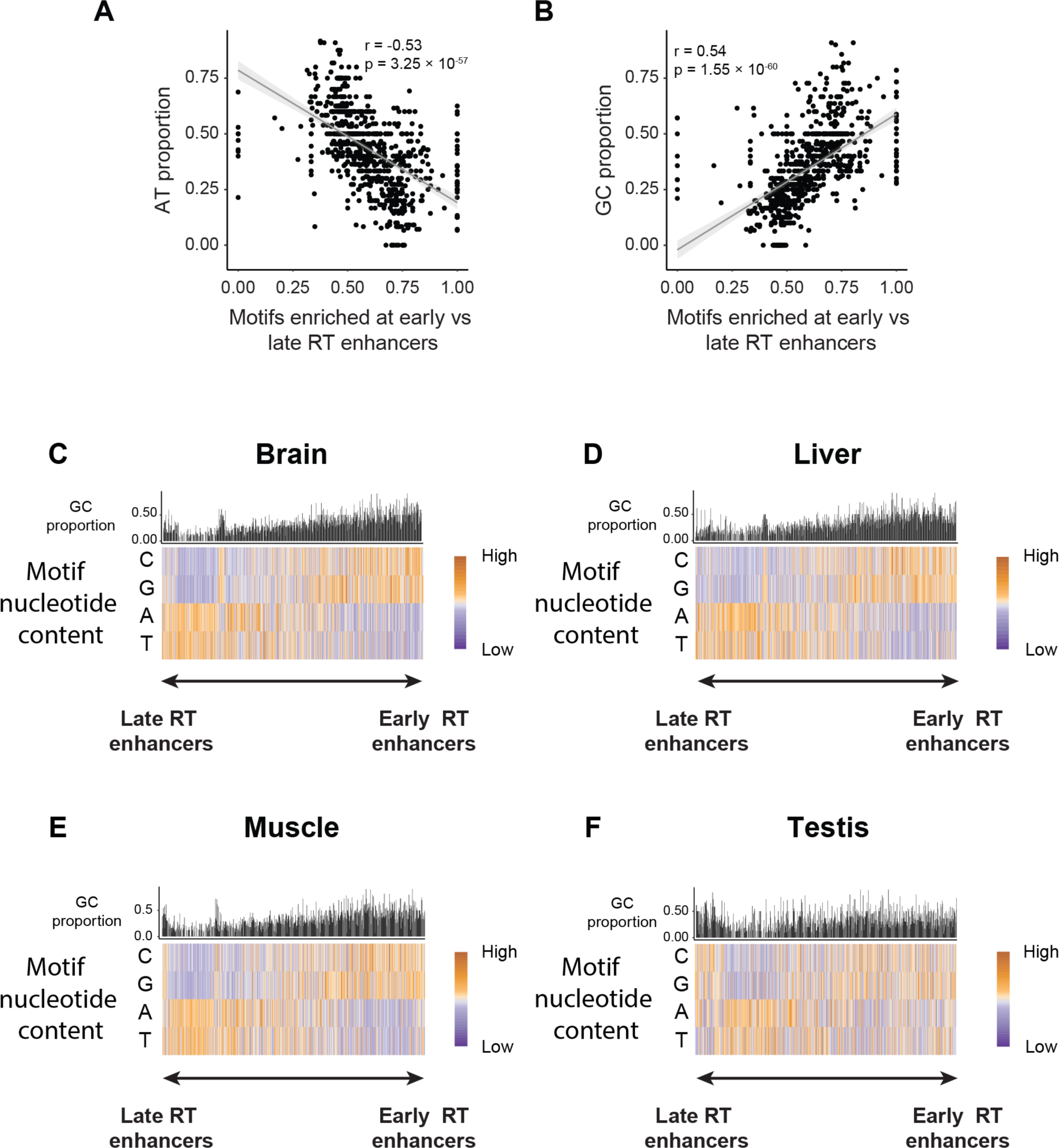
Nucleotide composition analyses at mouse enhancers based on DNA replication time. (**A-B)** Correlation between AT (**A)** and GC (**B)** proportion of TF binding motifs enriched at early vs late replicating mouse enhancers. Pearson’s r and p-value are indicated in each case. All tissues’ enhancers were included (brain, liver, muscle and testis). (**C-F)** Nucleotide composition of TF binding motifs enriched at early and late replicating mouse enhancers by tissue: (**C)** brain, (**D)** liver, (**E)** muscle and (**F)** testis (n = 6082, 6240, 2994 and 5094 brain, liver, muscle and testis enhancers, respectively; equal numbers of early RT and late RT enhancers for each tissue).

**Figure S15.**
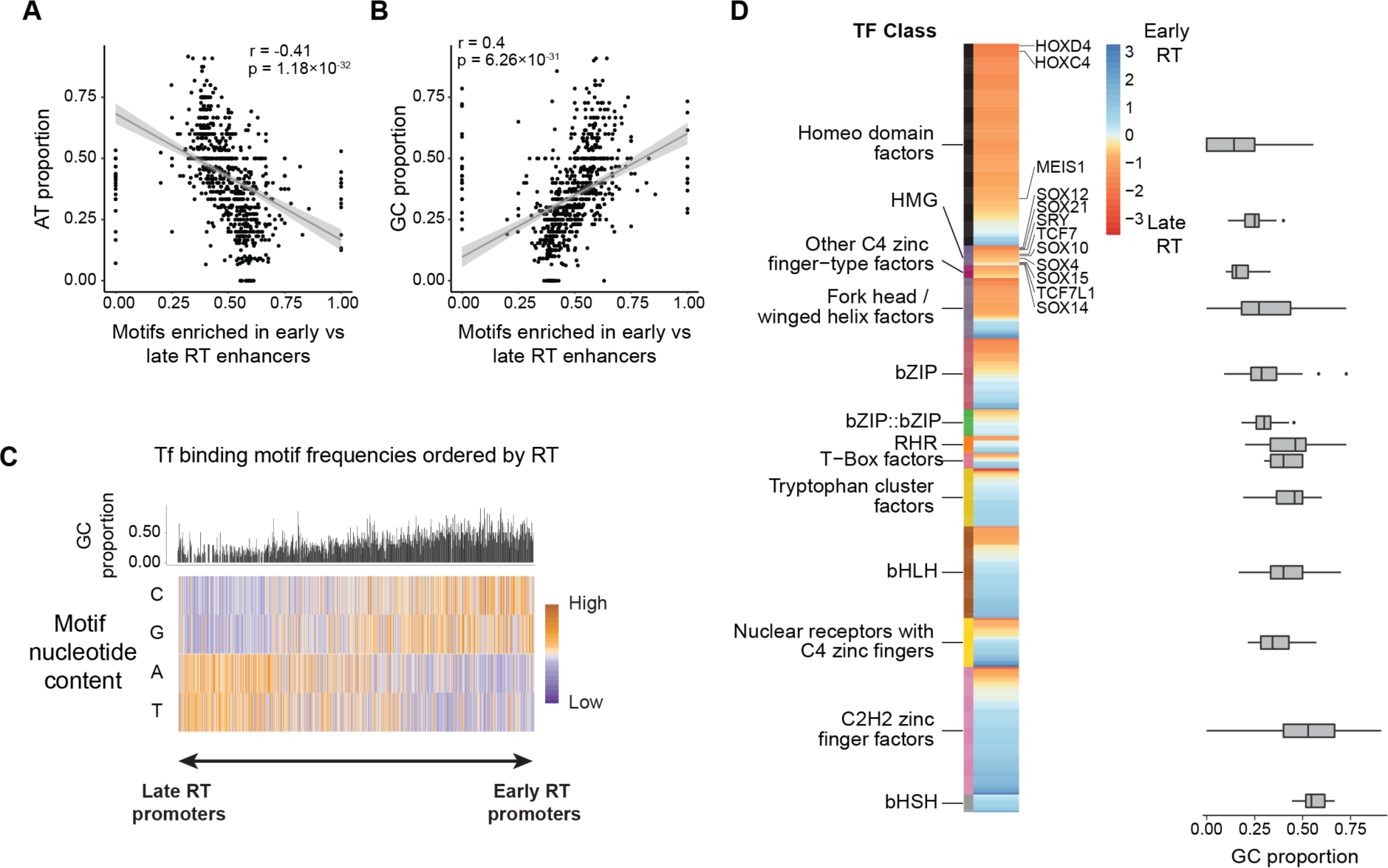
Nucleotide composition analyses at human liver promoters based on DNA replication time. (**A-B)** Correlation between AT (**A)** and GC (**B)** proportion of TF binding sites and TF relative enrichment at early vs late replicating promoters. Pearson correlation r and p-value are shown in each case. (**C)** Nucleotide proportion of TF binding motifs at early and late replicating human liver promoters (n = 2131 early RT enhancers and 2131 late RT enhancers). (**D)** Relative enrichment of TF binding motifs at early against late replicating promoters grouped by TF Class (left), some example motifs are indicated. The GC content of the motifs belonging to every TF Class is shown on the right (HMG = High-mobility group domain factors; bZIP = Basic leucine zipper factors; RHR = Rel homology region factors; bHLH = Basic helix-loop-helix factors; bHSH = Basic helix-span-helix factors). Only TF Classes with more than ten TFs are shown.

**Figure S16.**
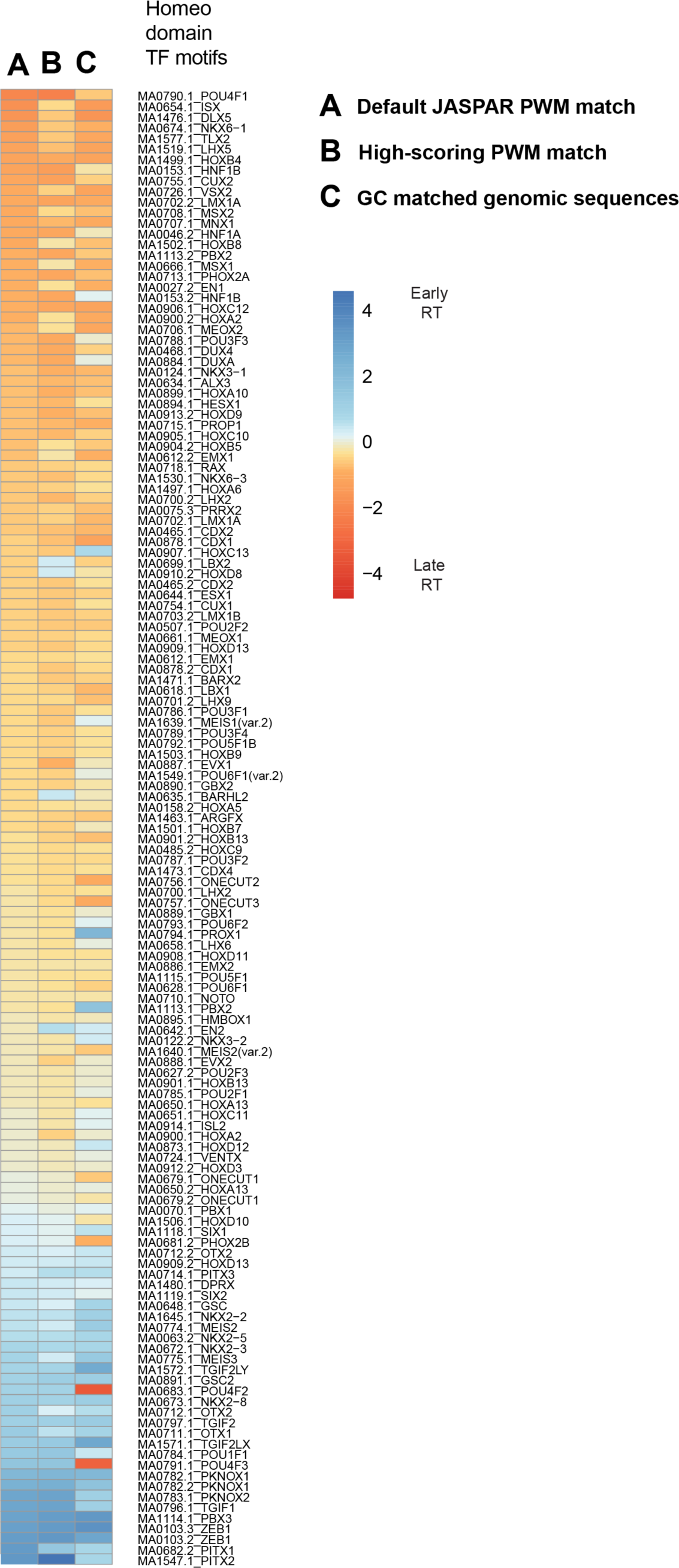
Relative enrichment of homeo domain factors at early and late replicating human liver enhancers. The left column of the heatmap shows the relative enrichment of homeodomain motifs at early and late replicating human liver enhancers. In the centre column only high scoring motifs are considered. The column on the right shows the relative enrichment of homeodomain factors in random regions of the genome matched by GC content. PWM IDs and transcription factor names are indicated. N = 139 homeo domain PWMs.

**Figure S17.**
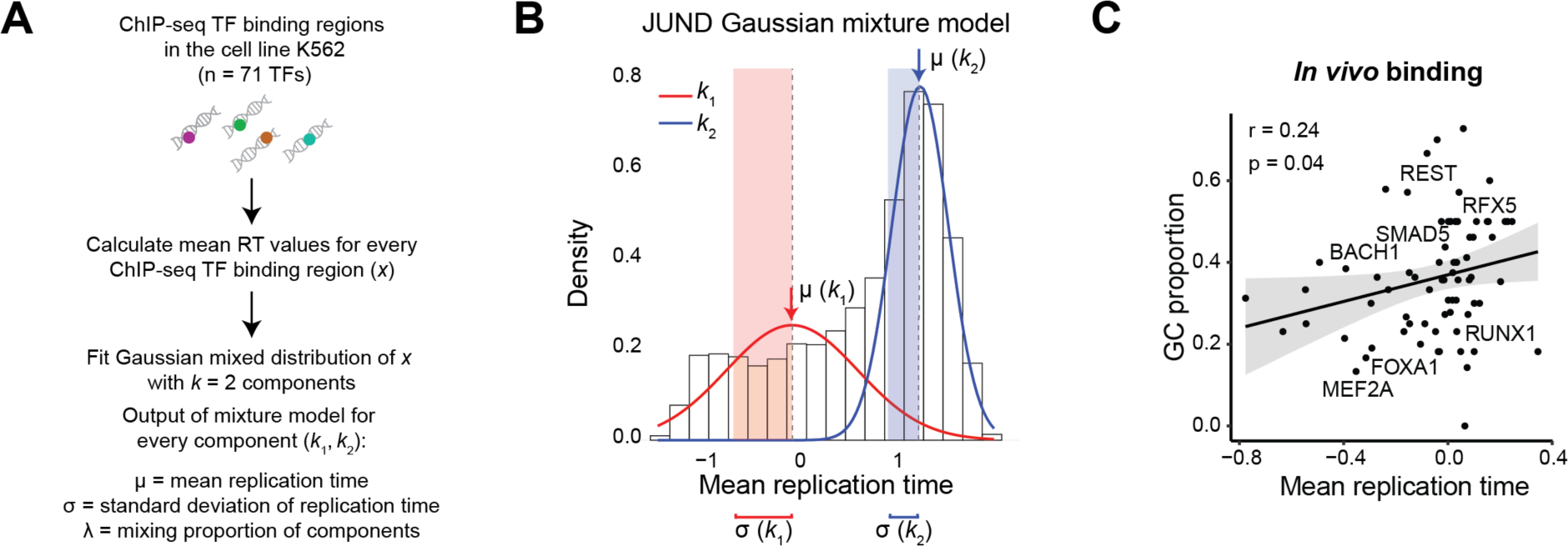
Gaussian mixture models of in vivo TF binding data in the cell line K562. (**A)** Pipeline of Gaussian mixture models. A model with two components (*k*_1_, *k*_2_) was fitted for the binding sites’ mean replication time of every TF (n = 71 TFs). After fitting a model, we got mean RT (μ), RT standard deviation (σ) and mixing proportion (λ) values for every component. (**B)** Example of mixture model (JUND). The distribution of JUND binding sites’ mean replication time is shown with a histogram. The distribution of the components *k*_1_ and *k*_2_ according to the fitted mixture model are represented with a red and a blue line, respectively. The mean RT value of every distribution is noted with a colour matched arrow. Shaded areas represent one standard deviation from the mean RT values. (**C)** Scatterplot of DNA replication time versus GC content of ChIP-seq binding sites in the cell line K562 for 71 TF ChIP-seq datasets. Mean replication time of binding sites in the later replicating cluster for each TF is shown (**Methods**). Pearson correlation coefficient (r) and p-value are shown. Shaded region represents the 95% confidence interval of the line of best fit.

**Figure S18.**
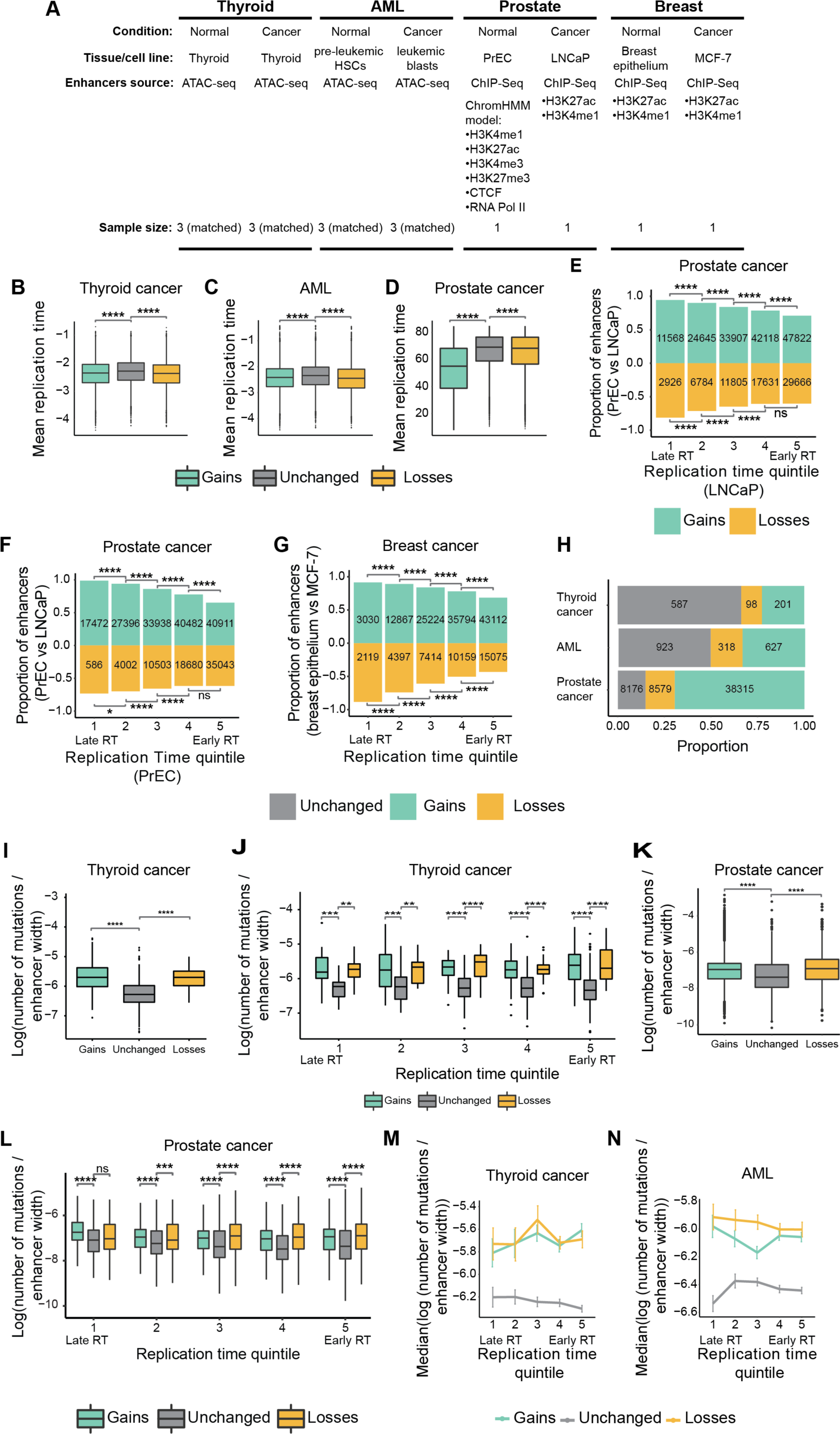
Enhancer turnover in cancer is associated with DNA replication time. (**A)** Summary of the datasets used to define enhancers in normal cell types and tissues and their cancer counterparts. The number of samples is indicated. (**B-D)** Mean replication time values for gains, losses and unchanged enhancers in thyroid cancer (**B),** AML (**C)** and prostate cancer (**D).** (Mann-Whitney *U*-test). (**E-G)** Proportion of gains and losses in prostate (based on LNCaP or PrEC replication time, **E** and **F**, respectively), and breast cancer (**G)** across replication time quintiles (Fisher’s exact test, alternative = “greater”, compared to unchanged enhancers). The number of enhancers is indicated in each case. (**H)** Proportion of mutations in gains, losses, and unchanged enhancers per cancer type. The number of mutations is displayed in each case. (**I)** Log transformed number of mutations normalized by enhancer width in thyroid cancer (Mann-Whitney *U*-test). (**J)** Log transformed number of mutations normalized by enhancer width in thyroid cancer across replication time quintiles (Mann-Whitney *U*-test**). (K)** Plot as (**I)** for prostate cancer. (**L)** Log transformed number of mutations normalized by enhancer width in prostate gains, losses, and unchanged enhancers across replication time quintiles (Mann-Whitney *U*-test). (**M-N)** Median log-transformed number of mutations normalized by enhancer width in thyroid cancer (**M)** and AML (**N)** gains, losses, and unchanged enhancers across replication time quintiles. Error bars represent standard error. The replication time shown for thyroid and AML was predicted from ATAC-seq data (predicted from thyroid and pre-leukemic HSCs, respectively). Significance notation: ‘ns’ P > 0.05; ‘***’ 1 ’ 10^−4^ < P ≤ 1 ’ 10^−3^; ‘****’ P ≤ 1 ’ 10^−4^. The number of enhancers and mutations in each cancer type can be found in **Supplemental Table S5**.

**Figure S19.**
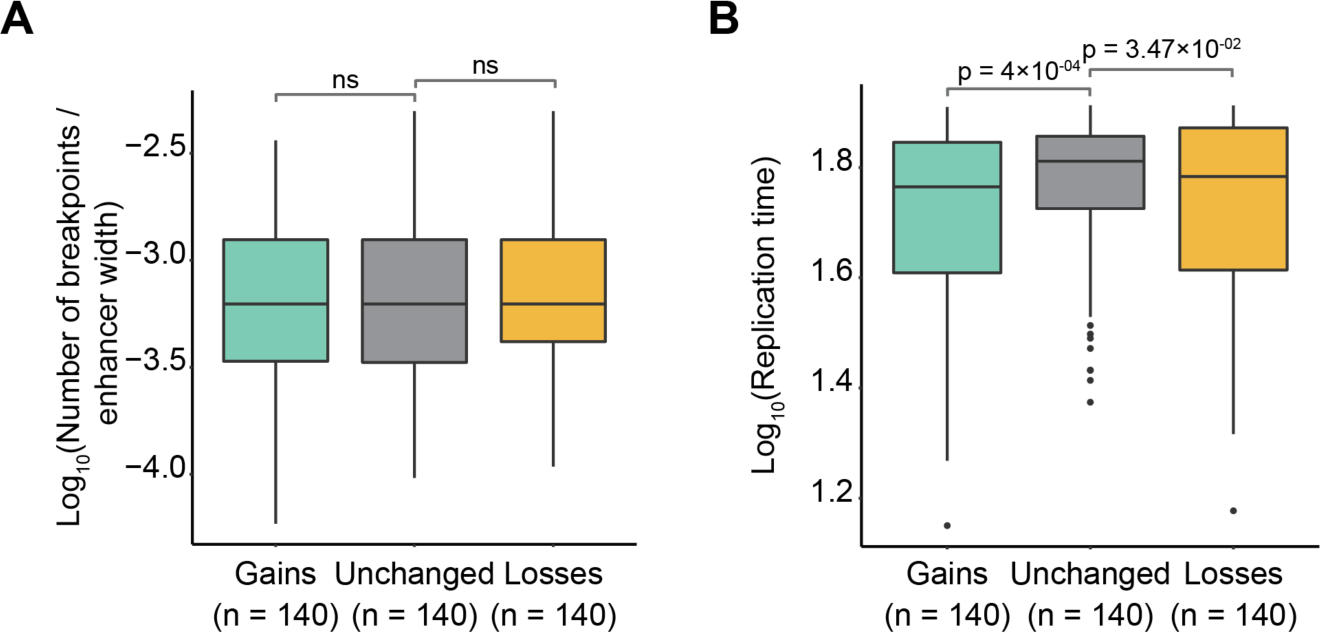
Replication time in prostate enhancers matched to recombination breakpoints. (**A)** Number of recombination breakpoints is matched after normalizing by enhancer width for gained, unchanged, and lost enhancers. Number of enhancers is indicated (Mann-Whitney *U*-test, significance code ‘ns’ p > 0.05). (**B)** Log_10_ transformed mean replication time of the enhancers shown in (**A)** (Mann-Whitney *U*-test, unchanged enhancers vs. gains or losses, alternative = “greater,” p-value is indicated for every comparison).

**Figure S20.**
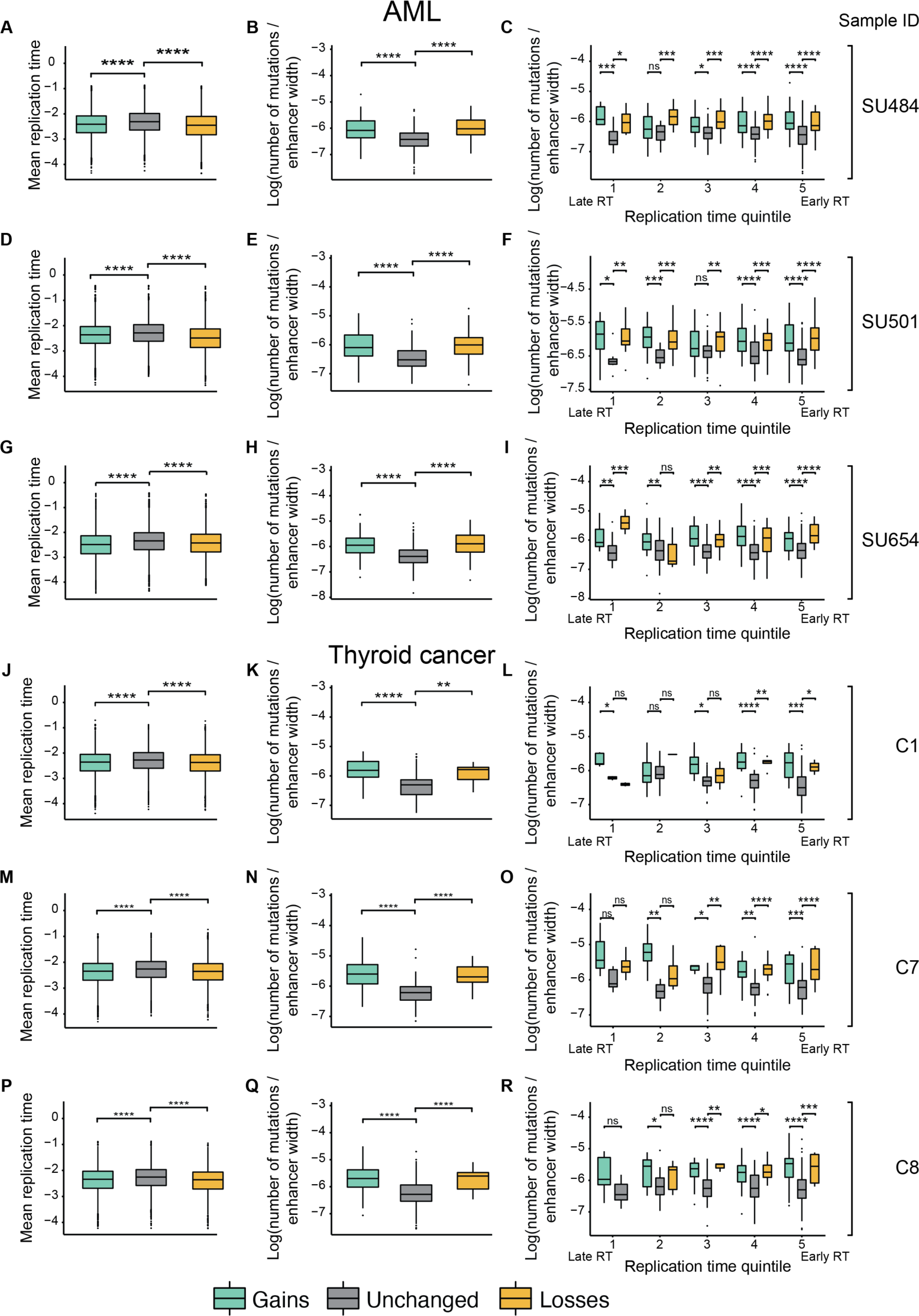
Enhancer mutations in individual cancer samples. Mean replication time and number of mutations in AML (**A-I)** and thyroid cancer (**J-R)** gains, losses and unchanged enhancers (n = 3 AML and n = 3 thyroid cancer patient samples). The first column of plots shows the mean replication time of gains, losses, and unchanged enhancers in individual AML and thyroid cancer patient samples (sample ID indicated on the right of the figure) (Mann-Whitney *U*-test). Second column shows the log-transformed number of mutations normalized by enhancer width in AML and thyroid cancer enhancers (Mann-Whitney *U*-test). Column 3 indicated the log transformed number of mutations normalized by enhancer width across replication time quintiles (Mann-Whitney *U*-test). ‘ns’ denotes p > 0.05, ‘****’ P ≤ 0.0001. The number of enhancers and mutations are indicated in **Supplemental Table S4**.

**Figure S21.**
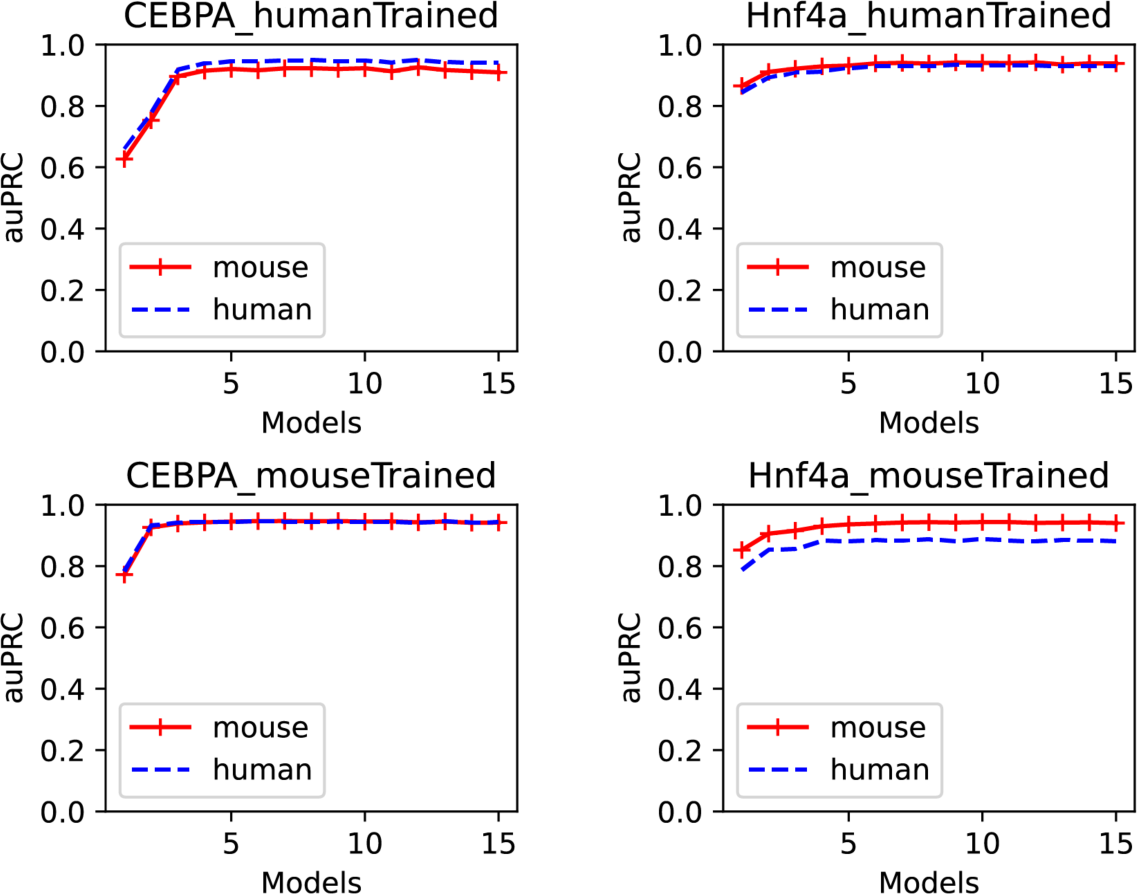
Model performance on validation datasets. The figure shows area under the Precision Recall curve (auPRC) values for models predicting CEBPA or HNF4A binding sites on the validation sets. Results from the models trained on human and mouse data are shown on the top and bottom rows, respectively. Model numbers are on the X-axis.

**Figure S22.**
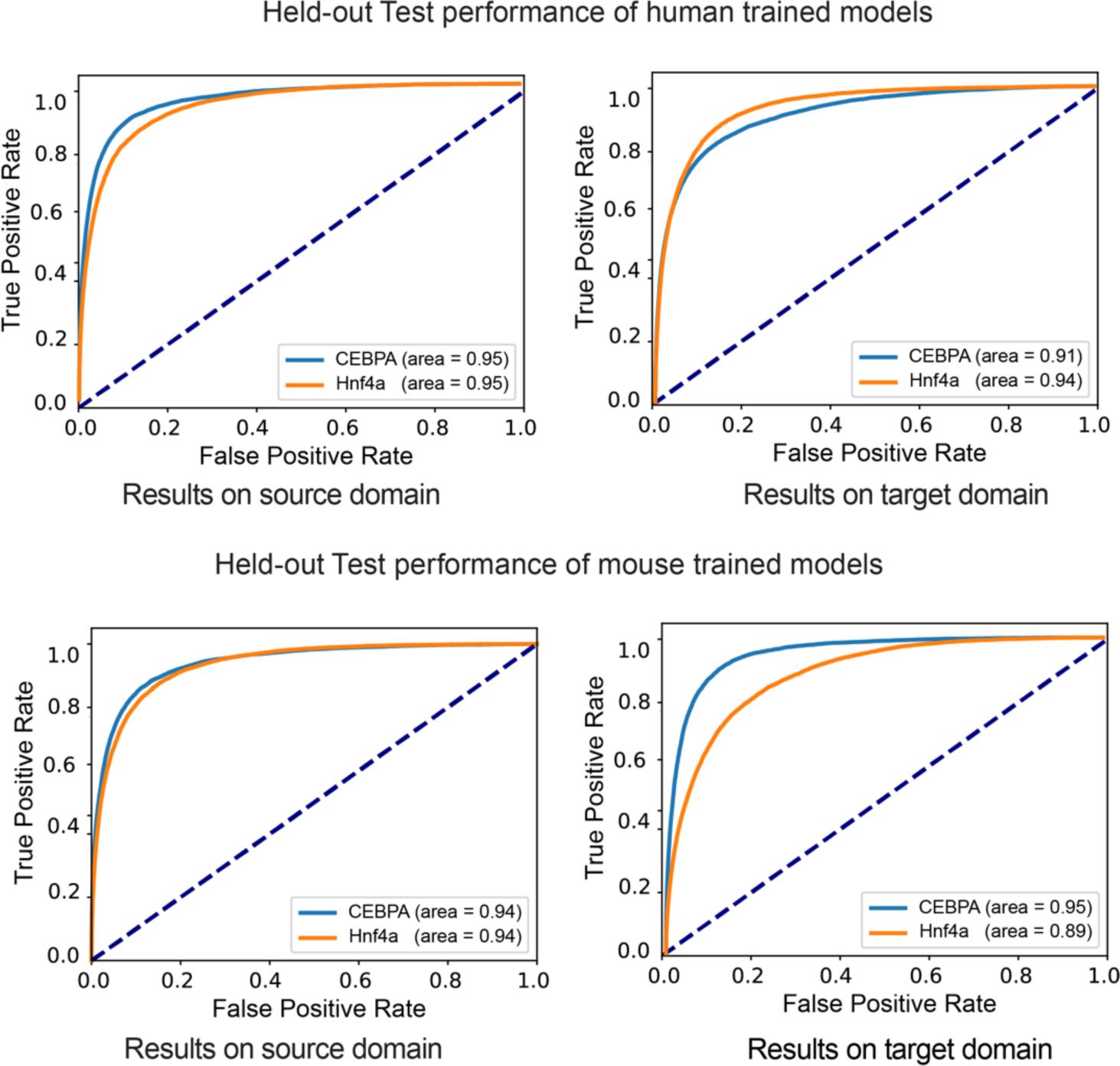
Held-out Test performance of the trained models. Receiver Operating Characteristic (ROC) curves depict the performance of models trained on human (top) and mouse data (bottom). In the plots, ROC curves for CEBPA and Hnf4a are shown in blue and orange, respectively. The corresponding areas under the ROC curve values are indicated. Predictions made on the same species are labelled as “source domain”, while “target domain” denotes the performance on the data from the other species (mouse or human).

## Supplemental Tables

**Table S1.** Fisher test for the differential overlap of recent and conserved enhancers with accessible genomic regions.

| Species | Tissue | N<br>conserved overlap<br>accessible regions | N<br>conserved do not<br>overlap accessible<br>regions | N<br>recent overlap<br>accessible regions | N<br>recent do not<br>overlap accessible<br>regions | P-value | Odds<br>ratio | Accession |
| --- | --- | --- | --- | --- | --- | --- | --- | --- |
| **Human** | Liver | 5119 | 8210 | 4451 | 5983 | 1 | 0.89 | GSE170971 |
| **Mouse** | Brain | 3204 | 33944 | 3144 | 19166 | 1 | 0.58 | ENCSR000COF |
| **Mouse** | Liver | 2392 | 26755 | 4838 | 19958 | 1 | 0.37 | ENCSR000CNI |
| **Mouse** | Muscle | 745 | 35315 | 861 | 21756 | 1 | 0.53 | ENCSR000CNX |

**Table S2.** Gaussian Mixture model’s output of the DNA replication time of TF binding sites in the cell line K562. ENCODE Accession numbers of the 71 TF binding datasets (ChIP-seq) used, TF IDs are also indicated. mu1 and mu2 represent the mean replication time values of the two Gaussian components generated with each TF binding dataset. Likewise, sd1 and sd2 represent the standard deviation of the components, l1 and l2 represent the components’ mixing proportions. and max1 and max2 represent the maximum value in every component.

| ENCODE accession | TF | mu1 | mu2 | sd1 | sd2 | l1 | l2 | mode1 | mode2 |
| --- | --- | --- | --- | --- | --- | --- | --- | --- | --- |
| ENCFF002CVN | ATF3 | -0.55 | 1.15 | 0.39 | 0.37 | 0.17 | 0.83 | 0.18 | 0.88 |
| ENCFF019PEL | ELK1 | 0.23 | 1.3 | 0.66 | 0.25 | 0.31 | 0.69 | 0.19 | 1.11 |
| ENCFF024TJO | YY1 | 0.04 | 1.25 | 0.68 | 0.28 | 0.35 | 0.65 | 0.21 | 0.91 |
| ENCFF043YZF | LEF1 | -0.16 | 1.2 | 0.6 | 0.31 | 0.37 | 0.63 | 0.25 | 0.8 |
| ENCFF076YZO | ETS1 | 0.25 | 1.27 | 0.69 | 0.26 | 0.32 | 0.68 | 0.19 | 1.04 |
| ENCFF085HTY | CTCF | -0.24 | 1.21 | 0.6 | 0.33 | 0.43 | 0.57 | 0.28 | 0.7 |
| ENCFF092TVM | NFIC | -0.13 | 1.25 | 0.67 | 0.29 | 0.46 | 0.54 | 0.27 | 0.73 |
| ENCFF106DAY | E2F1 | -0.03 | 1.26 | 0.68 | 0.29 | 0.43 | 0.57 | 0.25 | 0.79 |
| ENCFF113PMT | NFYB | 0.01 | 1.25 | 0.71 | 0.28 | 0.4 | 0.6 | 0.23 | 0.84 |
| ENCFF114IWY | ZNF143 | -0.01 | 1.26 | 0.68 | 0.28 | 0.41 | 0.59 | 0.24 | 0.85 |
| ENCFF144PPR | NRF1 | 0.06 | 1.25 | 0.68 | 0.28 | 0.36 | 0.64 | 0.21 | 0.91 |
| ENCFF150VTD | USF2 | -0.02 | 1.27 | 0.69 | 0.29 | 0.36 | 0.64 | 0.21 | 0.88 |
| ENCFF163VUK | ZKSCAN1 | 0.1 | 1.27 | 0.69 | 0.28 | 0.37 | 0.63 | 0.22 | 0.9 |
| ENCFF173TXA | GATA2 | -0.04 | 1.28 | 0.69 | 0.28 | 0.45 | 0.55 | 0.26 | 0.79 |
| ENCFF175IIE | NR2F1 | 0 | 1.27 | 0.67 | 0.28 | 0.4 | 0.6 | 0.24 | 0.85 |
| ENCFF175VSS | EGR1 | 0.04 | 1.25 | 0.69 | 0.28 | 0.37 | 0.63 | 0.21 | 0.89 |
| ENCFF178MOP | SMAD5 | -0.03 | 1.23 | 0.67 | 0.28 | 0.29 | 0.71 | 0.18 | 0.99 |
| ENCFF213EPU | ESRRA | 0.03 | 1.27 | 0.68 | 0.28 | 0.36 | 0.64 | 0.21 | 0.9 |
| ENCFF213EYD | JUND | -0.05 | 1.26 | 0.67 | 0.29 | 0.43 | 0.57 | 0.25 | 0.78 |
| ENCFF245LRG | ZBTB7A | 0.17 | 1.25 | 0.67 | 0.27 | 0.26 | 0.74 | 0.16 | 1.08 |
| ENCFF249EZR | TCF7 | 0.1 | 1.26 | 0.69 | 0.26 | 0.4 | 0.6 | 0.23 | 0.91 |
| ENCFF253FON | ZBED1 | 0.08 | 1.25 | 0.67 | 0.26 | 0.36 | 0.64 | 0.21 | 0.98 |
| ENCFF255EOB | NR2F2 | -0.01 | 1.26 | 0.67 | 0.28 | 0.4 | 0.6 | 0.24 | 0.84 |
| ENCFF290ESJ | REST | -0.16 | 1.24 | 0.63 | 0.3 | 0.44 | 0.56 | 0.28 | 0.74 |
| ENCFF290HTJ | STAT1 | -0.27 | 1.22 | 0.56 | 0.31 | 0.35 | 0.65 | 0.25 | 0.83 |
| ENCFF300XUA | SP1 | 0.16 | 1.26 | 0.68 | 0.27 | 0.28 | 0.72 | 0.17 | 1.06 |
| ENCFF302CPE | NFYA | 0.09 | 1.28 | 0.7 | 0.26 | 0.38 | 0.62 | 0.22 | 0.95 |
| ENCFF308IXJ | MAFF | -0.54 | 1.09 | 0.45 | 0.41 | 0.42 | 0.58 | 0.37 | 0.56 |
| ENCFF312RFN | TCF7L2 | 0.07 | 1.29 | 0.69 | 0.26 | 0.37 | 0.63 | 0.22 | 0.95 |
| ENCFF321KQD | CEBPB | -0.3 | 1.2 | 0.58 | 0.33 | 0.45 | 0.55 | 0.31 | 0.66 |
| ENCFF334FMW | USF1 | -0.02 | 1.24 | 0.67 | 0.28 | 0.41 | 0.59 | 0.24 | 0.83 |
| ENCFF384YDT | IRF2 | 0.01 | 1.27 | 0.68 | 0.28 | 0.42 | 0.58 | 0.24 | 0.83 |
| ENCFF392MUM | ELF1 | 0.08 | 1.26 | 0.67 | 0.28 | 0.38 | 0.62 | 0.23 | 0.88 |
| ENCFF408FQC | ZBTB33 | 0.02 | 1.24 | 0.66 | 0.29 | 0.32 | 0.68 | 0.19 | 0.95 |
| ENCFF417DTI | E2F6 | 0.1 | 1.24 | 0.68 | 0.28 | 0.33 | 0.67 | 0.19 | 0.95 |
| **ENCFF422BXE** | RUNX1 | 0.08 | 1.24 | 0.7 | 0.27 | 0.38 | 0.62 | 0.22 | 0.91 |
| **ENCFF423EMU** | BACH1 | -0.39 | 1.17 | 0.54 | 0.35 | 0.36 | 0.64 | 0.26 | 0.73 |
| **ENCFF426DUB** | JUNB | 0.03 | 1.28 | 0.7 | 0.26 | 0.46 | 0.54 | 0.26 | 0.83 |
| **ENCFF473GCH** | FOSL1 | -0.17 | 1.24 | 0.62 | 0.31 | 0.37 | 0.63 | 0.24 | 0.8 |
| **ENCFF492GXZ** | FOXK2 | 0.05 | 1.27 | 0.68 | 0.27 | 0.4 | 0.6 | 0.24 | 0.87 |
| **ENCFF497XOD** | MITF | -0.23 | 1.22 | 0.63 | 0.31 | 0.46 | 0.54 | 0.29 | 0.68 |
| **ENCFF502KHR** | ELF4 | 0 | 1.26 | 0.68 | 0.28 | 0.39 | 0.61 | 0.23 | 0.87 |
| **ENCFF517YCC** | CUX1 | -0.11 | 1.25 | 0.65 | 0.29 | 0.44 | 0.56 | 0.27 | 0.78 |
| **ENCFF544XKC** | PKNOX1 | -0.07 | 1.26 | 0.67 | 0.29 | 0.44 | 0.56 | 0.26 | 0.77 |
| **ENCFF547MLB** | TEAD4 | -0.15 | 1.24 | 0.64 | 0.3 | 0.43 | 0.57 | 0.27 | 0.75 |
| **ENCFF577UJR** | MEF2A | -0.35 | 1.18 | 0.6 | 0.32 | 0.36 | 0.64 | 0.24 | 0.8 |
| **ENCFF581YKY** | NEUROD1 | 0.02 | 1.26 | 0.68 | 0.28 | 0.35 | 0.65 | 0.21 | 0.92 |
| **ENCFF581ZZT** | ZNF282 | 0.07 | 1.25 | 0.67 | 0.27 | 0.33 | 0.67 | 0.2 | 0.99 |
| **ENCFF583CIY** | NR3C1 | 0.2 | 1.27 | 0.67 | 0.25 | 0.33 | 0.67 | 0.19 | 1.1 |
| **ENCFF613RNG** | MEIS2 | -0.15 | 1.23 | 0.64 | 0.31 | 0.45 | 0.55 | 0.28 | 0.71 |
| **ENCFF659WGE** | GATA1 | -0.03 | 1.28 | 0.69 | 0.28 | 0.41 | 0.59 | 0.24 | 0.84 |
| **ENCFF664XPS** | SPI1 | -0.09 | 1.25 | 0.67 | 0.3 | 0.43 | 0.57 | 0.26 | 0.76 |
| **ENCFF676NPW** | HES1 | -0.04 | 1.26 | 0.68 | 0.28 | 0.43 | 0.57 | 0.25 | 0.82 |
| **ENCFF676YJE** | MYC | 0.03 | 1.26 | 0.68 | 0.29 | 0.38 | 0.62 | 0.23 | 0.86 |
| **ENCFF706SJZ** | ZNF384 | 0.06 | 1.27 | 0.69 | 0.27 | 0.39 | 0.61 | 0.23 | 0.89 |
| **ENCFF710IEF** | ATF4 | -0.4 | 1.16 | 0.54 | 0.35 | 0.44 | 0.56 | 0.32 | 0.63 |
| **ENCFF716LRI** | RFX5 | 0.15 | 1.25 | 0.7 | 0.28 | 0.38 | 0.62 | 0.22 | 0.89 |
| **ENCFF730RUF** | IRF1 | -0.29 | 1.18 | 0.54 | 0.32 | 0.38 | 0.62 | 0.28 | 0.78 |
| **ENCFF765NAN** | FOXA1 | -0.31 | 1.2 | 0.5 | 0.35 | 0.37 | 0.63 | 0.3 | 0.72 |
| **ENCFF799HIG** | MAX | 0.03 | 1.26 | 0.67 | 0.28 | 0.36 | 0.64 | 0.21 | 0.91 |
| **ENCFF812QPN** | MAFK | -0.49 | 1.12 | 0.49 | 0.39 | 0.42 | 0.58 | 0.34 | 0.6 |
| **ENCFF833FCO** | GABPA | 0.04 | 1.26 | 0.68 | 0.28 | 0.36 | 0.64 | 0.21 | 0.92 |
| **ENCFF836LBG** | ZNF274 | -0.78 | 0.82 | 0.25 | 0.5 | 0.3 | 0.7 | 0.47 | 0.56 |
| **ENCFF891MVM** | ZNF263 | 0.11 | 1.26 | 0.7 | 0.27 | 0.36 | 0.64 | 0.2 | 0.96 |
| **ENCFF925DJG** | E2F8 | 0.15 | 1.28 | 0.7 | 0.26 | 0.35 | 0.65 | 0.2 | 1 |
| **ENCFF948TXN** | CREM | 0.02 | 1.26 | 0.69 | 0.29 | 0.39 | 0.61 | 0.23 | 0.85 |
| **ENCFF962FQM** | CTCFL | -0.08 | 1.24 | 0.67 | 0.3 | 0.35 | 0.65 | 0.21 | 0.87 |
| **ENCFF968KBN** | ATF2 | -0.63 | 1.02 | 0.4 | 0.46 | 0.41 | 0.59 | 0.41 | 0.52 |
| **ENCFF970LCB** | MXI1 | 0.12 | 1.27 | 0.66 | 0.27 | 0.34 | 0.66 | 0.2 | 0.98 |
| **ENCFF985TSH** | SREBF1 | 0.22 | 1.29 | 0.65 | 0.26 | 0.24 | 0.76 | 0.15 | 1.18 |
| **ENCFF990CFV** | POU5F1 | 0.35 | 1.32 | 0.66 | 0.23 | 0.33 | 0.67 | 0.2 | 1.14 |

**Table S3.** Number of enhancers and mutations per cancer type.

| **Cancer type** | **Sample** | **Enhancer type** | **Total number of enhancers** | **Number of mutations** |
| --- | --- | --- | --- | --- |
| **Prostate** |  | Gain | 159821 | 38315 |
| **Prostate** |  | Unchanged | 40742 | 8176 |
| **Prostate** |  | Loss | 68984 | 8579 |
| **Breast** |  | Gain | 119844 | NA |
| **Breast** |  | Unchanged | 36413 | NA |
| **Breast** |  | Loss | 39114 | NA |
| **Thyroid** | C1 | Gain | 14689 | 44 |
| **Thyroid** | C1 | Unchanged | 19014 | 118 |
| **Thyroid** | C1 | Loss | 15138 | 13 |
| **Thyroid** | C7 | Gain | 11142 | 36 |
| **Thyroid** | C7 | Unchanged | 16928 | 132 |
| **Thyroid** | C7 | Loss | 18665 | 63 |
| **Thyroid** | C8 | Gain | 22378 | 114 |
| **Thyroid** | C8 | Unchanged | 18915 | 307 |
| **Thyroid** | C8 | Loss | 7163 | 21 |
| **Thyroid** | Union | Gain | 41061 | 201 |
| **Thyroid** | Union | Unchanged | 28437 | 587 |
| **Thyroid** | Union | Loss | 34241 | 98 |
| **AML** | SU484 | Gain | 25297 | 185 |
| **AML** | SU484 | Unchanged | 18532 | 222 |
| **AML** | SU484 | Loss | 28276 | 111 |
| **AML** | SU501 | Gain | 34337 | 266 |
| **AML** | SU501 | Unchanged | 24174 | 234 |
| **AML** | SU501 | Loss | 31127 | 139 |
| **AML** | SU654 | Gain | 27277 | 141 |
| **AML** | SU654 | Unchanged | 40412 | 420 |
| **AML** | SU654 | Loss | 16300 | 53 |
| **AML** | Union | Gain | 58875 | 627 |
| **AML** | Union | Unchanged | 45184 | 923 |
| **AML** | Union | Loss | 48249 | 318 |

**Table S4.** Mean human and mouse enhancers width. The mean width and frequency of human and mouse enhancers is shown. Values are separated by enhancer type, mark and tissue where enhancers are functional.

| **Species** | **Class** | **Tissue** | **Mark** | **N enhancers** | **Mean width (bp)** |
| --- | --- | --- | --- | --- | --- |
| **Human** | Conserved | Liver | Active | 13329 | 3632 |
| **Human** | Recent | Liver | Active | 10434 | 3072 |
| **Mouse** | Conserved | Liver | Active | 13255 | 2977 |
| **Mouse** | Recent | Liver | Active | 8283 | 2029 |
| **Mouse** | Conserved | Liver | Poised | 15892 | 1223 |
| **Mouse** | Recent | Liver | Poised | 16513 | 785 |
| **Mouse** | Conserved | Brain | Active | 19146 | 2585 |
| **Mouse** | Recent | Brain | Active | 8804 | 2034 |
| **Mouse** | Conserved | Brain | Poised | 18002 | 1229 |
| **Mouse** | Recent | Brain | Poised | 13506 | 877 |
| **Mouse** | Conserved | Muscle | Active | 18616 | 2815 |
| **Mouse** | Recent | Muscle | Active | 8947 | 2154 |
| **Mouse** | Conserved | Muscle | Poised | 17444 | 1284 |
| **Mouse** | Recent | Muscle | Poised | 13670 | 880 |
| **Mouse** | Conserved | Testis | Active | 8381 | 2899 |
| **Mouse** | Recent | Testis | Active | 5584 | 1698 |
| **Mouse** | Conserved | Testis | Poised | 14365 | 1201 |
| **Mouse** | Recent | Testis | Poised | 18626 | 768 |

**Table S5.** Number of human liver enhancers alignable and with activity conservation in other species. Number of human liver enhancers with conserved activity in other species, aligned without conserved activity and not alignable. Species symbols are defined as in **Fig. S1**.

| **Class** | **Type** | **Species comparison** | **N enhancers** |
| --- | --- | --- | --- |
| **Conserved** | Align with no activity | Bbor | 1407 |
| **Conserved** | Align with no activity | Btau | 5719 |
| **Conserved** | Align with no activity | Cfam | 6807 |
| **Conserved** | Align with no activity | Cjac | 4198 |
| **Conserved** | Align with no activity | Cpor | 1742 |
| **Conserved** | Align with no activity | Csab | 5747 |
| **Conserved** | Align with no activity | Ddel | 1604 |
| **Conserved** | Align with no activity | Fcat | 6194 |
| **Conserved** | Align with no activity | Mbid | 1956 |
| **Conserved** | Align with no activity | Mdom | 2026 |
| **Conserved** | Align with no activity | Mfur | 1879 |
| **Conserved** | Align with no activity | Mmul | 5071 |
| **Conserved** | Align with no activity | Mmus | 6160 |
| **Conserved** | Align with no activity | Ocun | 6338 |
| **Conserved** | Align with no activity | Rnor | 5434 |
| **Conserved** | Align with no activity | Shar | 2772 |
| **Conserved** | Align with no activity | Sscr | 4422 |
| **Conserved** | Align with no activity | Tbel | 1643 |
| **Conserved** | Conserved activity | Bbor | 1579 |
| **Conserved** | Conserved activity | Btau | 5408 |
| **Conserved** | Conserved activity | Cfam | 4615 |
| **Conserved** | Conserved activity | Cjac | 5393 |
| **Conserved** | Conserved activity | Cpor | 1321 |
| **Conserved** | Conserved activity | Csab | 5753 |
| **Conserved** | Conserved activity | Ddel | 1382 |
| **Conserved** | Conserved activity | Fcat | 6603 |
| **Conserved** | Conserved activity | Mbid | 1030 |
| **Conserved** | Conserved activity | Mdom | 838 |
| **Conserved** | Conserved activity | Mfur | 1256 |
| **Conserved** | Conserved activity | Mmul | 6429 |
| **Conserved** | Conserved activity | Mmus | 3797 |
| **Conserved** | Conserved activity | Ocun | 3740 |
| **Conserved** | Conserved activity | Rnor | 4073 |
| **Conserved** | Conserved activity | Shar | 874 |
| **Conserved** | Conserved activity | Sscr | 4504 |
| **Conserved** | Conserved activity | Tbel | 803 |
| **Conserved** | Do not align | Bbor | 10343 |
| **Conserved** | Do not align | Btau | 2202 |
| **Conserved** | Do not align | Cfam | 1907 |
| **Conserved** | Do not align | Cjac | 3738 |
| **Conserved** | Do not align | Cpor | 10266 |
| **Conserved** | Do not align | Csab | 1829 |
| **Conserved** | Do not align | Ddel | 10343 |
| **Conserved** | Do not align | Fcat | 532 |
| **Conserved** | Do not align | Mbid | 10343 |
| **Conserved** | Do not align | Mdom | 10465 |
| **Conserved** | Do not align | Mfur | 10194 |
| **Conserved** | Do not align | Mmul | 1829 |
| **Conserved** | Do not align | Mmus | 3372 |
| **Conserved** | Do not align | Ocun | 3251 |
| **Conserved** | Do not align | Rnor | 3822 |
| **Conserved** | Do not align | Shar | 9683 |
| **Conserved** | Do not align | Sscr | 4403 |
| **Conserved** | Do not align | Tbel | 10883 |
| **Recent** | Align with no activity | Bbor | 1877 |
| **Recent** | Align with no activity | Btau | 4882 |
| **Recent** | Align with no activity | Cfam | 5277 |
| **Recent** | Align with no activity | Cjac | 3959 |
| **Recent** | Align with no activity | Cpor | 2034 |
| **Recent** | Align with no activity | Csab | 5430 |
| **Recent** | Align with no activity | Ddel | 1877 |
| **Recent** | Align with no activity | Fcat | 6759 |
| **Recent** | Align with no activity | Mbid | 1877 |
| **Recent** | Align with no activity | Mdom | 1915 |
| **Recent** | Align with no activity | Mfur | 2111 |
| **Recent** | Align with no activity | Mmul | 5430 |
| **Recent** | Align with no activity | Mmus | 4051 |
| **Recent** | Align with no activity | Ocun | 4408 |
| **Recent** | Align with no activity | Rnor | 3809 |
| **Recent** | Align with no activity | Shar | 2354 |
| **Recent** | Align with no activity | Sscr | 3758 |
| **Recent** | Align with no activity | Tbel | 1808 |
| **Recent** | Do not align | Bbor | 8557 |
| **Recent** | Do not align | Btau | 5552 |
| **Recent** | Do not align | Cfam | 5157 |
| **Recent** | Do not align | Cjac | 6475 |
| **Recent** | Do not align | Cpor | 8400 |
| **Recent** | Do not align | Csab | 5004 |
| **Recent** | Do not align | Ddel | 8557 |
| **Recent** | Do not align | Fcat | 3675 |
| **Recent** | Do not align | Mbid | 8557 |
| **Recent** | Do not align | Mdom | 8519 |
| **Recent** | Do not align | Mfur | 8323 |
| **Recent** | Do not align | Mmul | 5004 |
| **Recent** | Do not align | Mmus | 6383 |
| **Recent** | Do not align | Ocun | 6026 |
| **Recent** | Do not align | Rnor | 6625 |
| **Recent** | Do not align | Shar | 8080 |
| **Recent** | Do not align | Sscr | 6676 |
| **Recent** | Do not align | Tbel | 8626 |

